# A cross-scale assessment of productivity-diversity relationships

**DOI:** 10.1101/769232

**Authors:** Dylan Craven, Masha T. van der Sande, Carsten Meyer, Katharina Gerstner, Joanne M. Bennett, Darren P. Giling, Jes Hines, Helen R. P. Phillips, Felix May, Katherine H. Bannar-Martin, Jonathan M. Chase, Petr Keil

## Abstract

**Aim:** Biodiversity and ecosystem productivity vary across the globe and considerable effort has been made to describe their relationships. Biodiversity-ecosystem functioning research has traditionally focused on how experimentally controlled species richness affects net primary productivity (S→NPP) at small spatial grains. In contrast, the influence of productivity on richness (NPP→S) has been explored at many grains in naturally assembled communities. Mismatches in spatial scale between approaches have fostered debate about the strength and direction of biodiversity-productivity relationships. Here we examine the direction and strength of productivity’s influence on diversity (NPP→S) and of diversity’s influence on productivity (S→NPP), and how this varies across spatial grains.

**Location:** contiguous USA

**Time period:** 1999 - 2015

**Major taxa studied:** woody species (angiosperms and gymnosperms)

**Methods:** Using data from North American forests at grains from local (672 m^2^) to coarse spatial units (median area = 35,677 km^2^), we assess relationships between diversity and productivity using structural equation and random forest models, while accounting for variation in climate, environmental heterogeneity, management, and forest age.

**Results:** We show that relationships between S and NPP strengthen with spatial grain. Within each grain, S→NPP and NPP→S have similar magnitudes, meaning that processes underlying S→NPP and NPP→S either operate simultaneously, or that one of them is real and the other is an artifact. At all spatial grains, S was one of the weakest predictors of forest productivity, which was largely driven by biomass, temperature, and forest management and age.

**Main conclusions:** We conclude that spatial grain mediates relationships between biodiversity and productivity in real-world ecosystems and that results supporting predictions from each approach (NPP→S and S→NPP) serve as an impetus for future studies testing underlying mechanisms. Productivity-diversity relationships emerge at multiple spatial grains, which should widen the focus of national and global policy and research to larger spatial grains.

## Introduction

One of the most prominent questions in ecology is how to describe relationships between biodiversity and ecosystem-level productivity (Currie, 1991; Rosenzweig, 1995; Mittelbach *et al*., 2001; Balvanera *et al*., 2006; Adler *et al*., 2011; Cardinale *et al*., 2011, 2012; Hooper *et al*., 2012; Naeem *et al*., 2012; Tilman *et al*., 2014). Two fields of research with different motives have tried to understand causality between these variables (Loreau *et al*., 2001). The first examines how biodiversity varies across space as a result of different levels of productivity created by environmental variation (e.g., temperature, precipitation), and has resulted in a voluminous literature on the shapes of the patterns and their potential underlying causality (Connell & Orias, 1964; Currie, 1991; Rosenzweig, 1995; Waide *et al*., 1999; Mittelbach *et al*., 2001; Adler *et al*., 2011; Fraser *et al*., 2015). The second aims to quantify changes in vital ecosystem functions such as productivity following anthropogenically-induced changes in diversity (Schulze & Mooney, 1993; Tilman, 1999; Cardinale *et al*., 2012; Isbell *et al*., 2017). As a result of the different perspectives on the direction of causality, there remains considerable debate and confusion surrounding the relationship between diversity and productivity (Grace *et al*., 2016), which is exacerbated by differing spatial grains at which studies are conducted (Whittaker, 2010; Cardinale *et al*., 2011).

Recently, there has been growing interest in assessing biodiversity ecosystem functioning (BEF) relationships in real-world, non-experimental ecosystems over large geographic extents, but likely due to logistical constraints, relationships are typically measured at local spatial grains (Liang *et al*., 2016; Duffy *et al*., 2017; van der Plas, 2019). Results suggest that the positive effect of species richness on productivity and other ecosystem functions can be as, or more, important than abiotic environmental drivers’ effects on productivity, suggesting that diversity-productivity relationships can be even stronger in real-world communities than in controlled experiments (Duffy *et al*., 2017). However, to fully understand the influence of diversity on productivity, and vice versa, it is critical to recognize that traditional bivariate analyses may underestimate the strength of these relationships by not accounting for the effects of spatial grain, as well as those of biomass, shading, macro-climate, and management (Loreau *et al*., 2001; Cardinale *et al*., 2009; Oberle *et al*., 2009; Grace *et al*., 2016).

The striking mismatch between the spatial grains of BEF experiments (cm2 to m2; Cardinale *et al*., 2011), observational studies of BEF (0.04 to1.0 ha; Chisholm *et al*., 2013; Liang *et al*., 2016), and macroecological diversity-productivity correlations (m2 to thousands of km2; Mittelbach *et al*., 2001; Hawkins *et al*., 2003; Field *et al*., 2009; Adler *et al*., 2011) further obscures comparisons between perspectives. However, there is a diverse array of theoretical expectations for grain dependency of the effects of productivity on diversity (NPP→S) and of diversity on productivity (S→NPP), which predict effects to either strengthen or weaken as the spatial grain increases (Table 1; Gonzalez *et al*., 2020). For example, spatial turnover of species that are functionally equivalent within the regional grain can offset low species richness at local grains, resulting in a strengthening of S→NPP with increasing spatial grain. The effects of NPP→S are also hypothesized to increase with spatial grain, because higher NPP is associated with greater heterogeneity at larger spatial grains, which enhances coexistence of more species at the regional grain. Moreover, other components of a community, such as biomass, can mediate relationships between productivity and diversity via their effects on competitive dominance (Grace *et al*., 2016). These theoretical expectations have been supported by observational data for the effects of productivity on diversity (Mittelbach *et al*., 2001; Chase & Leibold, 2002; Belmaker & Jetz, 2011). In the case of BEF relationships (i.e. S→NPP), there is also empirical and theoretical support for grain dependence, which comes from a restricted range of small spatial grains (Luo *et al*.; Chalcraft, 2013; Hao *et al*., 2018).

**Table 1.** Overview of hypotheses predicting grain dependence of relationships between net primary productivity (NPP) and species richness (S).

| No. | Direction | Mechanism of grain dependence | Weakens or strengthens towards coarse grain? | Reference |
| --- | --- | --- | --- | --- |
| I | NPP → S and S → NPP | Spatially asynchronous demographic stochasticity impacts small populations (or small grains) and averages out over large grains. | Both NPP → S and S → NPP strengthen towards coarse grains | (Lande *et al.*, 2003) |
| II | NPP → S | At larger grains, higher NPP is associated with increased heterogeneity and/or dissimilarity of local patches, allowing for greater regional coexistence. | NPP → S strengthens towards coarse grains | (Abrams, 1988; Wright *et al.*, 1993; Chase & Leibold, 2002) |
| III | NPP → S | A statistical interaction between NPP and grain in their effect on S emerges as a consequence of increasing occupancy with NPP. | NPP → S weakens towards coarse grains | (Storch *et al.*, 2005) |
| IV | NPP → S | At very large grains (thousands of km<sup>2</sup> and larger), high productivity increases occupancy and population size, thus increasing the probability of reproductive isolation and speciation | NPP → S strengthens towards coarse grains | (Jetz & Fine, 2012) |
| V | S → NPP | Stochastic sampling effects dominate at small grains, resource partitioning at larger grains ('spatial insurance'), and their relative magnitude determines the grain dependency. | Both strengthening or weakening possible | (Loreau *et al.*, 2003; Cardinale *et al.*, 2004) |
| VI | S → NPP | Functionally redundant species at intermediate or coarse grains can compensate for low richness at local grains. | S → NPP strengthens towards coarse grains | (Srivastava & Vellend, 2005) |
| VII | S → NPP | With incomplete compositional turn-over, proportional changes in larger-grain richness are always less than proportional changes in smaller-grain richness such that the explanatory power of richness on changes in functioning decreases with spatial scale. | S → NPP strengthens towards coarse grains until species richness saturates | (Thompson *et al.*, 2018) |

Here, we aim to address the dual nature by which productivity influences diversity (NPP→S) and diversity influences productivity (S→NPP) across spatial grains by combining structural equation models (SEM) and random forest models (RFs) to explicitly account for the bidirectionality of NPP→S and S→NPP. Using SEM, we propose and test hypothesis-based models (Fig. S1) that estimate the direction and strength of NPP→S and S→NPP. Next, we use RFs, an assumption-free machine learning approach (Breiman, 2001; Hastie *et al*., 2009), to quantify the relative importance of predictors of species richness and productivity. We examine both hypothesized directions of the relationship, along with a number of important covariates that influence both diversity and productivity, such as biomass, precipitation, temperature, and forest age, using a comprehensive observational dataset of North American forests at fine (area = 672 m^2^; *n =* 46,211 plots), medium (median area = 1,386 km^2^; *n =* 1,956 spatial units), and coarse spatial grains (median area = 35,677 km^2^; 98 spatial units). We specifically ask whether the influence of productivity on diversity (NPP→S) was stronger or weaker than the influence of diversity on productivity (S→NPP), and how these relationships manifest across grains in real-world ecosystems.

## Methods

### Data

#### Geographic extent and grain

We conducted analyses across the contiguous USA at three spatial grains (Fig. 1): (1) fine grain (46,211 plots, 672 m^2^ or 0.000672 km^2^ each), (2) intermediate grain (1,956 units, median 1,386 km^2^) created by aggregating US counties to larger units based on the forested area within them (see ‘spatial aggregation algorithm’ below), and (3) coarse grain (95 units, median 35,677 km^2^) created by further aggregating the intermediate grain units. We restricted our analyses to forested areas to make comparisons within and among spatial grains in similar ecosystems. For the intermediate and coarse grains, we defined an area as forested if it fell into a 1 km^2^ pixel with non-zero forest biomass following Blackard *et al*. (2008).

**Fig. 1.**
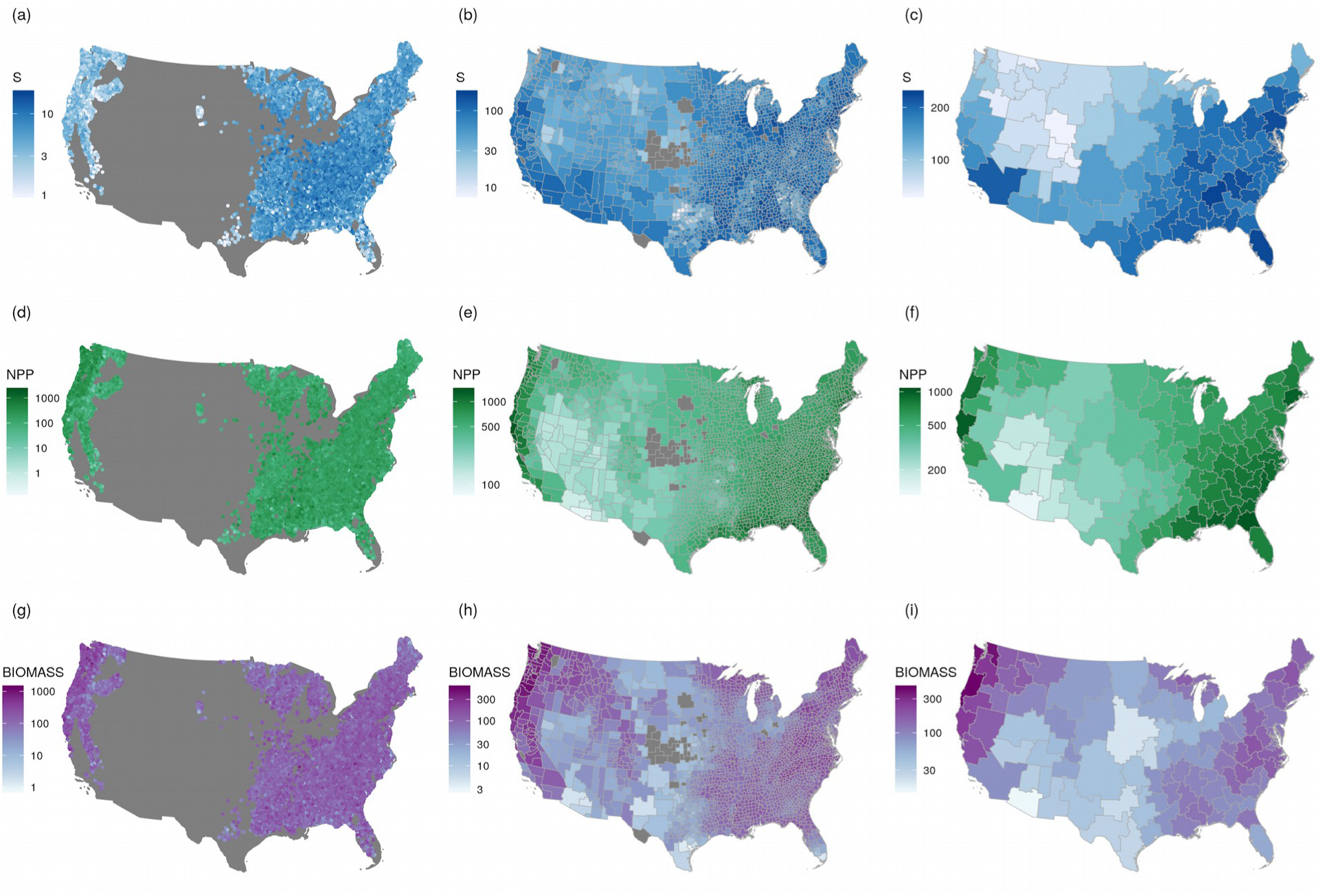
Maps of species richness (S), MODIS-derived net primary productivity (NPP) [gC/m^2^/year], and biomass [Mg/ha] of forests at three spatial grains across the contiguous USA. The values in all plots are on log_10_ scale.

#### Species richness (S)

For all spatial grains, we estimated diversity as species richness (S) because it is the most commonly used and best understood metric of biodiversity, although other measures of diversity may be better predictors of net primary productivity (Paquette & Messier, 2011; Cadotte, 2015; Venail *et al*., 2015). We extracted S at the fine spatial grain from the Forest Inventory and Analysis National Program (FIA) database v. 1.7.0 (USDA Forest Service, 2017). We restricted our analysis to plots on forested land that were sampled using the national FIA design (plot design code 1)(Burrill *et al*., 2018). All plots were surveyed between 1998 and 2016, each consisting of four circular 168 m^2^ sub-plots with a total area of 672 m^2^ ha in which all individuals larger than 12.7 cm diameter at 1.3 m were recorded and identified to species level. For each plot we pooled data from all subplots to estimate S. In total, our final dataset included 344 woody species and 93,771 plots. We estimated S at the intermediate and coarse spatial grains by counting the number of unique woody species in each spatial unit using data for the contiguous USA provided in The Biota of North America Program’s (BONAP) North American Plant Atlas (Kartesz, 2015).

#### Taxonomic harmonization of species names

We cleaned scientific names from the FIA and BONAP datasets and harmonized them to accepted species based on The Plant List (2013) and the Taxonomic Name Resolution Service (2018), following the protocol described in Meyer et al. (2016). We included hybrid forms, but excluded any names that could not be resolved to the species level.

#### Filtering of species occurrences

We restricted our analyses to tree species that likely occur in forests. At the fine spatial grain, we included native and alien species. At the intermediate and coarse spatial grains, however, we excluded alien species because we could not be certain if they occurred in forests as many are cultivated, particularly in urban ecosystems (Kowarik, 2008; Pearse *et al*., 2018). We therefore filtered the BONAP data to native species classified as ‘trees’ in BONAP’s taxonomic query database (Kartesz, 2015). We further filtered out 70 county-level occurrences of 5 non-woody species from the BONAP dataset. Species woodiness was inferred from woodiness data (Zanne *et al*., 2013) and species lists of trees, shrubs and subshrubs (USDA NRCS, 2018), except for 37 species without such data for which we instead inferred woodiness from online searches or assumed resemblance among congeneric species. We also filtered out 8 FIA plot-level species occurrences and 1,595 BONAP county-level species occurrences that we deemed unlikely to be forest occurrences, as inferred from independent species occurrences within forested pixels recorded in FIA plots and Global Biodiversity Information Facility (GBIF) point-occurrence records (Downloaded via https://www.gbif.org/ on 26th September 2016; doi:10.15468/dl.mka2y5; Supplementary Note 1). To make species richness data internally consistent across the different spatial grains, we added a further 6,593 quality-vetted county-level forest occurrences of woody species from FIA plot records to the 282,991 occurrences in the taxonomically harmonized BONAP dataset.

#### Net primary productivity (NPP)

At all spatial grains, we calculated NPP using MODIS-derived estimates, which we further supplemented with plot-derived estimates at the fine spatial grain. Briefly, we calculated NPP using the MODIS-derived MOD17 A3 product (Zhao *et al*., 2005; Zhao & Running, 2010), which gives annual values of NPP as gC m^-2^ yr^-1^ in 30 arc-sec pixels (roughly 1 km^2^ around the equator). Here, NPP is defined as the annual sum of daily net photosynthesis minus the cost of growth and maintenance of living cells in permanent woody tissue. We averaged the annual values from 2000 to 2015 for each pixel, and then averaged these across the intermediate and coarse grains. We use MODIS-derived NPP in the analyses presented in the main text to ensure comparability across spatial grains.

At the fine spatial grain, we also estimated NPP using plot-derived data. For a large subset of plots in the FIA database that have been measured at least twice between 1999 and 2015 (n = 46,211, on average plots re-measured every 5.8 years), we calculated net annual net aboveground C change (gC m^-2^ y^-1^). This was measured as the net change in aboveground tree C between two measurements as the sum of aboveground C growth of living trees, ingrowth by recruitment, and loss from tree mortality (NPPmort; Chen & Luo, 2015). Tree-level carbon was estimated by multiplying tree-level biomass (see below) by 0.48, but we recognize that gymnosperms may have higher carbon content than that of angiosperms (Thomas & Martin, 2012). For plots with more than two inventories, tree productivity was calculated for each period and then averaged. NPPmort was weakly correlated with MODIS-derived NPP at the fine spatial grain (*r* = 0.19), suggesting that it may capture different processes. Therefore, we provide the analyses using the plot-derived NPP at the fine spatial grain in the Supplementary Information. Importantly, results concerning the strength of the S-NPP relationship were qualitatively similar for both NPP measures.

#### Biomass (BIOMASS)

At all spatial grains, we derived biomass values using a map of aboveground forest biomass of the USA, which is derived by modeling FIA plot biomass as a function of geospatial predictor variables (Blackard *et al*., 2008). This data layer had a grain of 250 × 250 m^2^, therefore, the average within each of the intermediate- and coarse-grain spatial units was taken.

For analyses using plot-derived NPP, we estimated tree-level biomass at the fine spatial grain using generalized biomass equations developed for North American tree species (Chojnacky *et al*., 2013). For each FIA plot we calculated aboveground biomass (Mg ha^-1^) as the sum of individual biomass of living trees per hectare.

#### Number of trees (N)

At the fine scale, we estimated the number of trees directly from each FIA plot. For the intermediate and coarse spatial grains, we estimated the number of trees using a global map of tree density (Crowther *et al*., 2015). As the grain of the data layer was 1 × 1 km^2^, average tree density was calculated within each spatial unit at the intermediate and coarse spatial grains.

#### Forest age (AGE) and management (MANAGED)

For each plot in the fine-scale dataset, we extracted forest age and management history from the FIA data set. Forest age is estimated using dendrochronological records (Burrill *et al*., 2018). Management regime was a binary variable that indicated whether any forest management activity, e.g. harvest, thinning, tree planting, had been observed in any inventory or not.

At the intermediate and coarse grain, forest age was calculated as the average forest age from NASA NACP 1 km^2^ resolution layer (Pan *et al*., 2012). Management regime at the intermediate and coarse grains was calculated as the proportion of managed FIA plots within all FIA plots that were within each spatial unit.

#### Climatic variables

For all grains, we used WorldClim (Hijmans *et al*., 2005) bioclimatic variables at 30 sec resolution. Many of the WorldClim variables are strongly collinear with one another, or with other variables in the analysis (Table S1, Fig. S1). Thus, only three variables that captured different aspects of the climate were selected; mean annual temperature (BIO1; ANN.TEMP), mean precipitation (BIO12; ANN.PREC), and temperature seasonality (BIO4; TEMP.SEAS). At the fine scale, for each FIA plot we extracted the values of the 30 sec pixel in which the plot was found. For the intermediate and coarse grains, we averaged the values across all pixels within each spatial unit.

#### Elevation range (ELEV.RANGE)

We used elevation range as a proxy for topographic and habitat heterogeneity, a variable that has been shown to be a good predictor of species richness (Stein *et al*., 2014). The USGS SRTM1 dataset (USGS, 2009) with 1 sec (approx. 30 × 30 m^2^) resolution was used for all spatial grains. At the fine-scale, we calculated a 250 m diameter buffer around each FIA plot and calculated the elevation range using all 1 sec SRTM pixels within the buffer. At the intermediate and coarse scale, elevation range was calculated as the difference between the minimum and maximum elevation points within each spatial unit.

#### Species pools (S.POOL)

We calculated regional species pools for each spatial grain as probabilistic dispersal pools (Karger *et al*., 2016). For each intermediate-grain spatial unit and each species in our data set, we first estimated the species’ probability of being part of the unit’s species pool as the joint probabilities that dispersal might happen between that unit and any of the species’ intermediate-grain occurrences within the contiguous US. Due to insufficient data on species’ dispersal abilities, we assumed that dispersal probability between focal units and species’ occurrences would decay with great-circle distance between the respective regions’ centroids. We explored five alternative exponential distance-decay functions, with scaling coefficients *P* that determined the probability of a species occurring in neighboring units would disperse to the focal unit of 0.975, 0.95, 0.90, 0.80, and 0.60. We chose the function with *P* = 0.8, which exhibited the strongest correlation between species pool and species richness at all spatial grains (Fig. S2). Finally, we calculated species pools for each spatial unit as the sum of all species’ individual probabilities of dispersal from any of their respective occurrences. For each coarse-grain unit, we summed the species’ joint probabilities of dispersal between any of their intermediate-grain occurrences and any of the intermediate-grain units nested within the coarse unit. For fine-grain units, we assumed that their species pools would equal those of the intermediate-grain spatial units in which they were nested.

All of the variables used in our analyses are listed and summarized in Table S1 and visualized in Fig. S1.

#### Spatial aggregation algorithm

Because US counties vary dramatically in their area (Fig. S3), from Falls Church (VA) being as small as 5.1 km^2^, to San Bernardino (CA) with 52,109 km^2^, it is difficult to assign one categorical grain size to county-level data. Thus, we aggregated county data for species richness to create new spatial units, with the goal to minimize variation in forested area (*A*) between spatial units. We achieved this using a greedy algorithm which worked as follows: (1) Calculate variance (*V*_*1*_) of forested area (*A*) across all counties. (2) Randomly select a focal county with a probability proportional to 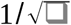, which will most likely select counties with small *A*. (3) Randomly choose a county adjacent to the focal county and merge it with the focal county. (4) Update the variance (*V*_*2*_) of forested area across all spatial units in the USA and compare it to the original variance *V*_*1*_. If the *V*_*2*_ < *V*_*1*_, the algorithm accepts the merged unit and returns to step one. If the variance does not decrease, the algorithm repeats step 3 until *V*_*2*_ < *V*_*1*_, with the maximum number of attempts of 1,000. If the variance still does not decrease even after 1,000 attempts, the algorithm rejects the merge, and returns to step one. The algorithm started with 3,107 counties, and we first terminated it when 1,956 merged spatial units were created. We classified these spatial units as the intermediate spatial grain (Fig. 1). We then allowed the algorithm to continue until it reached 98 merged spatial units, which we classified as the coarse spatial grain (Fig. 1). Although the algorithm substantially reduced variation in area within both spatial grains (Fig. S3), it did not eliminate the variation entirely. For this reason, we used area as a covariate in the statistical analyses at the intermediate and coarse spatial grains.

#### Stratified random sampling

Large areas of the contiguous US are environmentally homogeneous, while other parts are environmentally unique and small. We employed stratified random sampling (Cochran, 1977) for the fine and intermediate spatial grains in order to (1) enhance environmental representativeness of the data, (2) prevent excessive statistical leverage of the large number of data points from homogeneous areas and (3) reduce spatial pseudoreplication (autocorrelation) by increasing the geographic distance between data points. We first identified 11 strata at the fine and intermediate grains respectively, using multivariate regression trees with S, NPP and biomass as response variables and all covariates as predictors (Fig. 1). We then took a random and proportionally sized sample of spatial units from each strata (fine grain, N = 1,000; intermediate grain, N = 500). We did not use stratified random sampling at the coarse spatial grain because of the small number of spatial units (N = 98). The spatial locations of the stratified samples are in Fig. S4. All of the analyses presented here, as well as our main conclusions, are based on these stratified sub-samples of the data.

#### Data transformation and standardization

Prior to analysis, species richness, biomass, N, NPP, and area were natural-log transformed to meet normality assumptions of the standardised major-axis regressions and SEMs.

## Data Analyses

We quantified simple bivariate relationships between diversity and productivity for each spatial grain using standardised major-axis regression with the ‘sma’ function in the R package ‘smatr’ (Warton *et al*., 2012). We then used two complementary statistical approaches to assess the impacts of diversity and productivity and vice versa while simultaneously accounting for covariates that influence both.

First, we fitted structural equation models (SEMs), which allow the assessment of indirect effects including feedback loops, address causality, and take into account potential collinearity among covariates (Grace *et al*., 2010; Shipley, 2016). The paths in our candidate SEMs were based on previous evidence of causal links between S, biomass, and NPP (Fig. 2; Grace *et al*., 2016). Second, to better understand the relative importance of each variable in explaining variation in the response variables within models, we fitted random forest models (RFs) (Hastie *et al*., 2009). The results from SEMs provide insight into differences among models (i.e. between the two causal pathways per spatial grain, and among spatial grains), while results from RFs provide additional insights into the relative importance of different predictors variables within models.

**Fig. 2.**
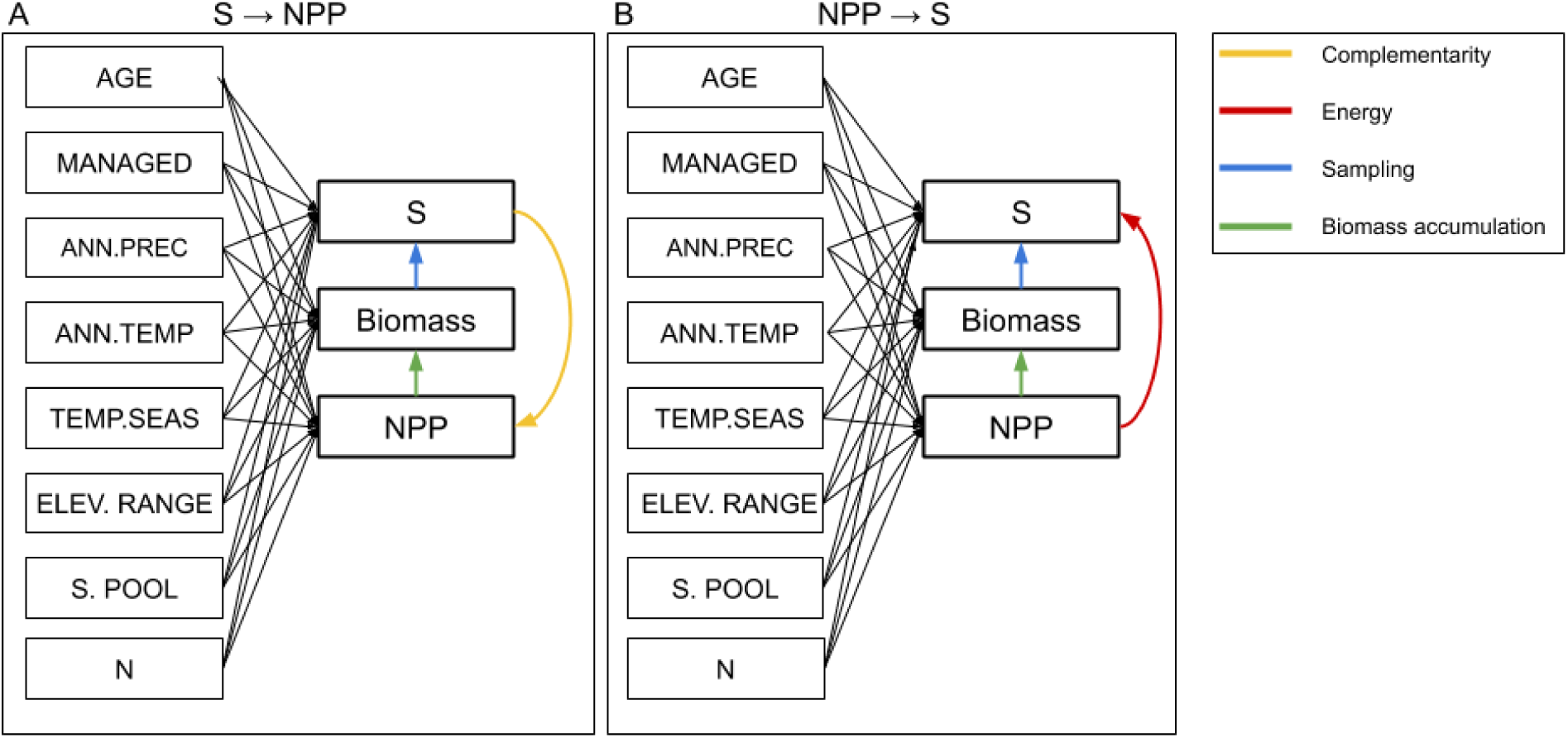
Hypothetical causal models for structural equation models (SEM) testing the relative importance of species richness (S) on net primary productivity (NPP) (‘S→NPP’; A) and NPP on S (‘NPP→S’; B) in forests across the contiguous USA at three spatial grains. Paths in color represent possible ecological mechanisms influencing the direction of the relationship; yellow paths represent complementarity effects, red paths represent ‘species-energy’ relationships, blue paths represent sampling (or niche) effects and green paths represent biomass accumulation. Black paths are relationships of additional covariates with S, NPP, and BIOMASS and are not hypothesized to occur in a particular direction. AGE is forest age, MANAGED is forest management, ANN.PREC is mean annual precipitation, ANN.TEMP is mean annual temperature, TEMP.SEAS is temperature seasonality, ELEV.RANGE is elevation range, S.POOL is the regional species pool, and N is the number of individuals. At the intermediate and coarse spatial grains, we added AREA to the SEMs to account for differences in the area of spatial units. S, BIOMASS, NPP, and AREA were natural log transformed prior to analysis.

### Structural Equation Modelling (SEM)

To test the relative importance of S on NPP (‘S→NPP’) and NPP on S (‘NPP→S’) we fitted two SEMs per spatial grain. For each SEM, we started with a ‘saturated’ model, which included the relationships between S, NPP, and biomass, and relationships of all additional covariates on S, NPP, and biomass (except for area at the fine spatial grain) (Fig. 2). The S→NPP model evaluated how S directly affects NPP and how NPP indirectly affects S via biomass and, therefore, included a feedback loop. The NPP→S model tested the direct effect of NPP on richness and, unlike the S→NPP model, did not include a feedback loop. This way, we tested the direct effect of S on NPP (S→NPP model), the direct effect of NPP on S (NPP→S), and the indirect effect of NPP on S (included in both models).

Model fit can only be tested on unsaturated models, i.e. those that have at least one missing path. Therefore, we removed the path with the lowest standardized path coefficient from the model. As SEMs had an equal number of paths, we could compare model fit across all models within each spatial grain using their unadjusted R^2^ values. After excluding the additional paths, path coefficients of S, NPP, and biomass remained qualitatively the same, and model fit to the data were still accepted (Chi-square test; P>0.05). This indicates that the models are identifiable and their results are robust. Therefore, we did not further reduce the model, and models maintained the same number of paths within each scale. Because models at the fine spatial grain including the number of individuals (N) did not fit the data well (P < 0.05), we excluded this variable. Models at the intermediate and coarse spatial grains including N fit the data well (P >0.05), but we present models without N for consistency with the fine spatial grain and because the sampling effects captured by N are also captured by area.

To assess the differences among scales in the relationships between S, NPP and biomass for each model, we compared the standardized regression coefficients using their 95% confidence intervals. All SEMs were fitted using the ‘sem’ function of the ‘lavaan’ package in R (Rosseel, 2012).

### Random forest models (RFs)

To assess the relative importance of each variable in predicting the response variables within models, we used random forest models (RFs) (Breiman, 2001; Liaw & Wiener, 2002; Hastie *et al*., 2009). We used the ‘randomForest’ function in the R package ‘randomForest’, with all RF models produced using the default settings: 500 trees, one third of predictors sampled in each tree, sampling with replacement of the entire dataset, and terminal node size of 5.

At each of the three spatial grains we fitted two RFs, one with S as a response variable and the other with NPP as a response variable. All predictors that were used in the SEMs were used in the RF models (including biomass). To quantify the relative importance of each predictor, we calculated the mean decrease of squared error across all 500 trees using the ‘importance’ function. The importance values were then scaled between 0 and 1, with 1 being the most important predictor. Using the function ‘partialPlot’, we extracted the partial responses of S and NPP to visualize the relationship between the two variables after accounting for all other covariates.

### Non-linear responses and spatial autocorrelation

SEMs offer the advantage of modelling complex, causal relationships (Grace *et al*., 2010; Shipley, 2016), but they can be difficult to fit to data with non-linear responses or spatial pseudoreplication. While it is possible to model non-linearity in SEMs, e.g. using polynomials (Grace *et al*., 2010; Shipley, 2016), this often comes at the cost of interpretability. A similar problem applies when it comes to another prevalent problem of observational geographic data: spatial autocorrelation, which statistical models have so far addressed by modelling it either in residuals, or in the response (Dormann *et al*., 2007). However, because of the causal loop in the SEMs (Fig. 2), the key response variables are also predictors, which prevented us from estimating spatial autocorrelation. In our analyses, we account for these issues in the following manner: (1) In the SEM analyses, we keep the relationships linear, given the approximately linear pairwise relationships between the raw NPP, S and biomass data (Figs. S5-7). (2) In the SEM analyses we do not directly model spatial autocorrelation. (3) We address spatial autocorrelation in the random forest analysis by allowing the algorithm to model smooth geographic trends in the response (by including the X and Y spatial coordinates as predictors), and we measure spatial autocorrelation in the response and in residuals. (4) We allow the random forest analysis to detect non-linear responses.

### Reproducibility

All data on species richness, biomass, NPP, covariates, and R code used for the data processing and analyses are available on Figshare (DOI: 10.6084/m9.figshare.5948155) under a CC-BY license.

## Results

Spatial patterns in productivity (NPP) and richness (S) emerged at coarser spatial grains, with higher S and NPP usually observed in the eastern USA than in the western USA (Fig. 1). Biomass, a time-integrated measure of NPP that also influences diversity, also exhibited similar patterns (Fig. 1). Bivariate relationships between S and NPP exhibited scale dependence (Fig. 3). While not significantly correlated at the fine spatial grain (standardised major axis regression: R^2^ = 0.00, P = 0.73), S and NPP were significantly correlated at the intermediate (standardised major axis regression: R^2^ = 0.15, P < 0.001) and coarse spatial grains (standardised major axis regression: R^2^ = 0.35, P < 0.001). The slope of S-NPP increased from 0.86 (95% confidence intervals: 0.80, 0.94) at the intermediate spatial grain to 1.23 (95% confidence intervals: 1.05, 1.45) at the coarse spatial grain. Similar patterns were observed when using plot-derived estimates of NPP at the fine spatial grain (Fig. S8).

**Fig. 3.**
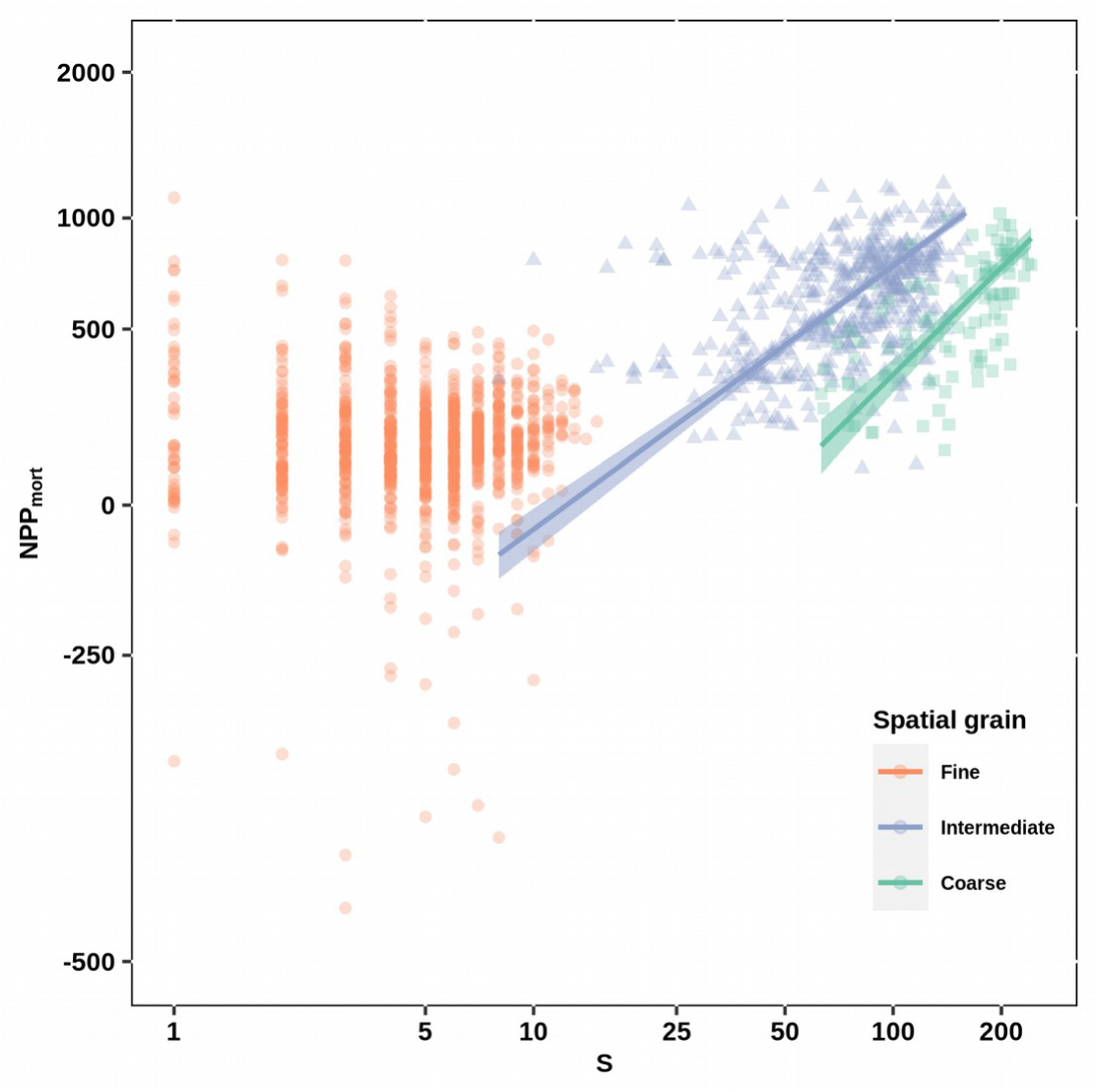
Bivariate relationships between observed species richness (S) and productivity (NPP) of forests at three spatial grains across the contiguous USA. Solid lines are standardised major-axis regressions fitted at each spatial grain and shaded areas are 95% confidence intervals; only regressions with statistically significant slopes (P<0.05) were visualised. NPP is MODIS-derived at all spatial grains. Note that axes are on the natural log scale. Analyses were performed using stratified random samples of 1000, 500 and 98 spatial units at the fine, intermediate and coarse spatial grains, respectively.

### Structural Equation Models (SEM)

We examined relationships between species richness and net primary productivity (NPP) across spatial grains using two SEMs for each spatial grain: the first (S→NPP) testing the direct effect of S on NPP and the indirect effect of NPP on S (via biomass), and the second (NPP→S) testing both the direct and indirect effects of NPP on S (Fig. 4). In both SEMs, environmental variables (e.g., mean annual precipitation, mean annual temperature, temperature seasonality, and elevation range), size of the species pool, forest age, and management were used to explain variation in S, biomass, and NPP. At the intermediate and coarse grains, we also included area (of each spatial unit) to account for variation in species richness due to sampling effects.

**Fig. 4.**
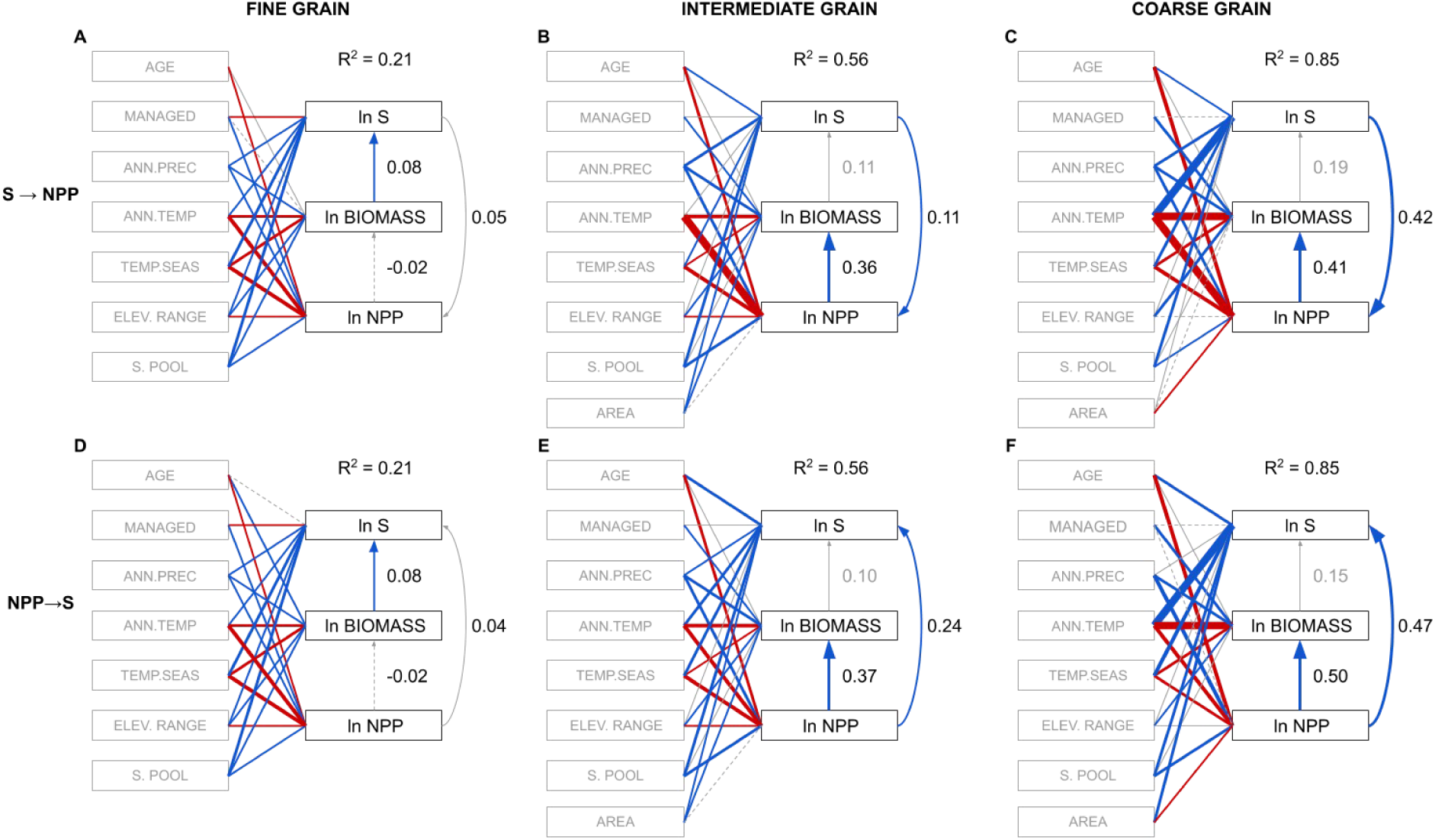
Structural equation models (SEM) testing the influence of diversity (S) on productivity (NPP)(‘S → NPP’; A, B, C) and that of NPP on S (‘NPP → S’; D, E, F), once controlling for environmental variables (e.g., mean annual precipitation, mean annual temperature, temperature seasonality, and elevation range), size of the species pool, forest age, and management, in forests across the contiguous USA at three spatial grains. All models fit the data well at all spatial grains (P-value of the Chi-square test > 0.1; Table S1). Boxes represent measured variables and arrows represent relationships among variables. Solid blue and red arrows represent significant (P< 0.05) positive and negative standardized path coefficients, respectively, and their width is scaled by the corresponding standardized path coefficient. Solid and dashed gray arrows represent non-significant (P>0.05) positive and negative standardized path coefficients, respectively. R^2^ is the average of R^2^ values for S, BIOMASS, and NPP. NPP is MODIS-derived at all spatial grains. AGE is forest age, MANAGED is forest management, ANN.PREC is mean annual precipitation, ANN.TEMP is mean annual temperature, TEMP.SEAS is temperature seasonality, ELEV.RANGE is elevation range, S.POOL is the regional species pool, and AREA is area. S, BIOMASS, NPP, and AREA were natural log transformed prior to analysis.

Both models fit the data well for all spatial grains (P-value of the Chi-square test > 0.1; Table S2). At each spatial grain, both SEMs had similar R^2^ values averaged over S, biomass and NPP, indicating a similar fit of the model to the data. R^2^ values for both SEMs increased with spatial grain, from 0.21 at the fine grain, to 0.56 at the intermediate grain and 0.85 at the coarse grain. Generally, the strength of effects of S → NPP and NPP → S were similar within each spatial grain, but both increased in strength with increasing spatial grain (Figs. 4 & 5). At the fine spatial grain, we found a weak direct effect of S → NPP (Fig. 4A) and NPP → S (Fig. 4D), and effectively a null indirect effect of NPP on S via biomass (standardized path coefficient of indirect effect = −0.002; Fig. 4A). At the intermediate spatial grain, we found a similarly strong direct effect of S on NPP (standardized path coefficient of direct effect = 0.11, Figs. 4B and 5) as NPP on S (standardized path coefficient of direct effect = 0.24; Figs. 4E and 5) and weak indirect effects of NPP on S via biomass (standardized path coefficient of indirect effect = 0.04; Fig. 4B). Similarly at the coarse spatial grain, we found strong direct effects of S on NPP (0.42, Fig. 4C and 5) and of NPP on S (0.47, Fig. 4F and 5) and weak indirect effects of NPP on S via biomass (standardized path coefficient of indirect effect = 0.08; Fig. 4C).

**Fig. 5.**
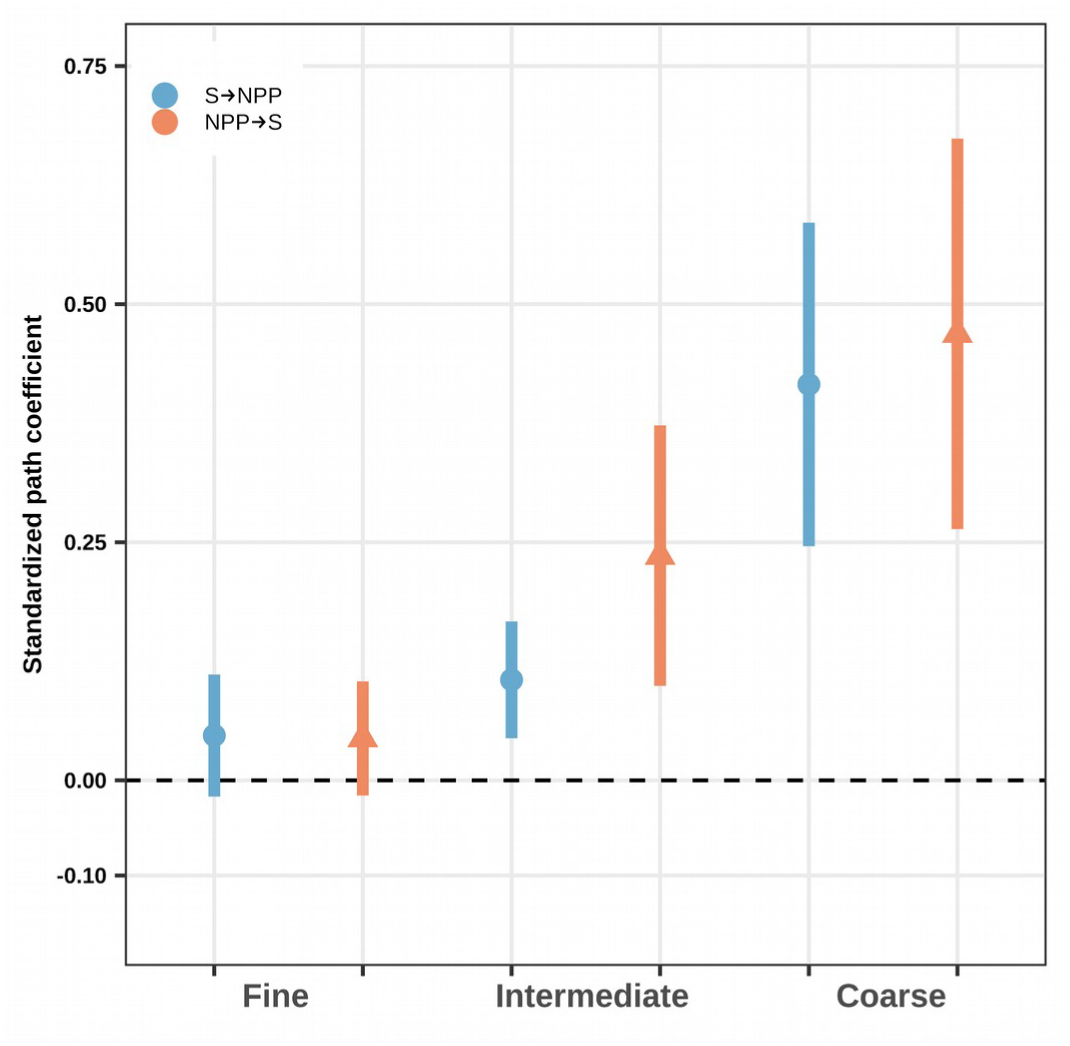
Direct effects of diversity on productivity (S → NPP) and productivity on diversity (NPP → S) estimated with structural equation models (SEM) in forests across the contiguous USA at three spatial grains. Points are standardized path coefficients and solid lines are 95% confidence intervals.

Overall, the SEMs suggest that the productivity-diversity relationship increases in strength with spatial grain, and both relationships (S→NPP and NPP→S) explain similar amounts of variation. At all spatial grains, our SEMs do not conclusively show stronger support for one direction of causality over the other. Similar patterns were observed when using plot-derived estimates of NPP (Fig. S9; Table S2), except for the direction of direct effects of S on NPP and NPP on S, which was negative.

### Random forest models (RFs)

To assess the relative importance of each predictor of species richness and NPP, and to provide an assumption-free alternative to the SEMs that also accounts for spatial autocorrelation, we fitted two random forest models for each of the three spatial grains: one with NPP and the other with S as response variables. We found that species richness was one of the weakest predictors of NPP relative to other predictors at all spatial grains (Fig. 6A), with management, forest age, MAP, and biomass being the most important predictors (Fig. 6A). The overall explained variation of NPP also increased from the fine to the two coarser spatial grains, from 0.64 at the fine spatial, to 0.89 at the intermediate spatial grain and 0.88 at the coarse spatial grain.

**Fig. 6.**
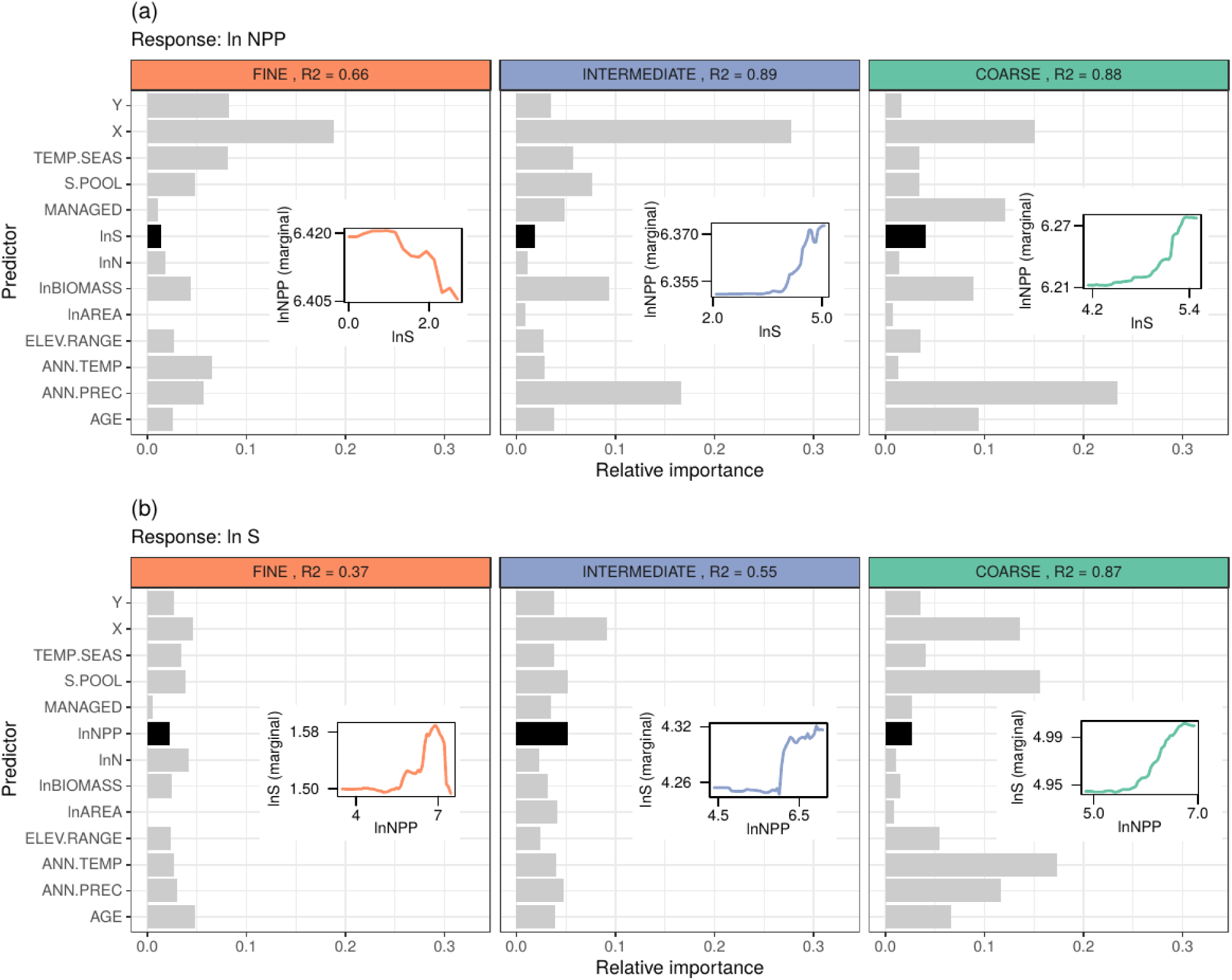
Relative variable importance from random forest models explaining (a) MODIS-derived NPP and (b) species richness (S) at three spatial grains. Relative variable importance is the mean decrease in squared error caused by each of the variables, rescaled such that it sums up to the total pseudo R^2^ of the whole random forest model. The curves in the insets show shapes of the marginal responses of ln NPP or ln S after accounting for all of the covariates. Y and X are latitudinal and longitudinal coordinates of the US National Atlas projection, TEMP.SEAS is temperature seasonality, S.POOL is the regional species pool, MANAGED is forest management, lnNPP is MODIS-derived NPP, lnN is the number of individuals, lnBIOMASS is biomass, lnAREA is area of the spatial unit, ELEV.RANGE is elevation range, ANN.TEMP is mean annual temperature, ANN.PREC is mean annual precipitation, and AGE is forest age. For explanation of variables see Table S1.

We found that NPP was an important predictor of S (with a positive effect) only at the intermediate spatial grain (Fig. 6B), but was less important relative to other predictors at fine and coarse spatial grains. For S, we found that species pool, mean annual temperature and precipitation, and forest age were the best predictors, and their importance increased towards coarse spatial grains (Fig. 6). In line with the SEM analyses, the overall explained variation of S increased towards coarse spatial grains, from 0.39 at the fine grain to 0.55 at the intermediate and 0.87 at coarse grains (see Fig. S10 for predicted vs. observed values).

In all RF analyses, there is a clear East-West spatial component in both S and NPP (represented by the X coordinate in Fig. 6), which was not explained by any of the other predictors. This spatial component was stronger for NPP than for S. Residual autocorrelation in all of the RF models was negligible (Fig S11). Finally, we also fitted all of the RFs with local plot-derived measures of productivity (as an alternative to the MODIS-derived productivity used in the main analyses), showing that the strength of the S-NPP relationships were similar across all NPP measures (Fig. S12).

## Discussion

The first important result is the similar magnitude of the S→NPP (Grace *et al*., 2016) and NPP→S (Mittelbach *et al*., 2001; Hawkins *et al*., 2003; Šímová *et al*., 2011) relationships at all grains. This reflects, in part, that both productivity and species richness have many environmental and geographical drivers in common (Lavers & Field, 2006), which complicates distinguishing correlation from causation, even when using SEMs (Grace *et al*., 2010; Shipley, 2016). There are two possible interpretations of this result: (i) it may indicate that diversity’s causal effects on productivity and productivity’s causal effects on diversity operate simultaneously, which was suggested by (Grace *et al*., 2016), but never demonstrated on observational data from large spatial grains. Alternatively (ii), if only one direction of the diversity-productivity relationship is real and causal, it may be possible to fit another model assuming the opposite direction because of multicollinearity in the data or non-identifiability of the causal direction (Petersen & van der Laan, 2014). Without large-grain experiments that manipulate diversity in ways that mimic biodiversity change (i.e. species gains and losses) in real-world ecosystems (Loreau *et al*., 2001; Wardle, 2016; Hillebrand *et al*., 2018; Manning *et al*., 2019; Gonzalez *et al*., 2020), we see little hope for resolving this with contemporary data and approaches.

Our second important result is that both S→NPP and NPP→S strengthen from the fine to the intermediate grain, and in the case of the SEM both relationships continue strengthening towards the coarsest grain. While grain-dependent shifts are often expected (Table 1), this had not been shown previously with empirical data for S→NPP using spatial grains coarser than several hectares (Luo *et al*.; Chisholm *et al*., 2013; Hao *et al*., 2018). If the S→NPP direction is the real causal one, then our results from SEM and RF analyses support several theoretical expectations (Table 1) and give further impetus to efforts quantifying biodiversity effects in naturally assembled ecosystems at broad spatial scales (Isbell *et al*., 2018). If the NPP→S direction is the real causal one, then our results are in line with (Lavers & Field, 2006; Field *et al*., 2009), but are in contrast with (Storch *et al*., 2005; Belmaker & Jetz, 2011), particularly when upscaling from the fine grain to intermediate grain, where both the SEM and RF analyses give congruent results. Intriguingly, a third possibility is that both NPP→S and S→NPP are real and that they operate simultaneously, as suggested by our SEM results. In this case, we are unaware of any theory that considers how this reciprocal relationship would be expected to change with increasing spatial grain. The one caveat applicable to interpreting any direction of diversity-productivity relationships is that of demographic stochasticity (mechanism I in Table1), which may weaken both NPP→S and S→NPP, or their synergistic interplay, at fine spatial grains. In our study, the strong local effect of demographic stochasticity appears plausible given the small area of the forest plots (672 m^2^) and small population sizes (12.24 ± 0.02 trees per plot; range = 1-157 trees per plot) therein. This would suggest that temporal changes in local scale biodiversity (Dornelas *et al*., 2014; Magurran *et al*., 2018) may have under-appreciated effects on ecosystem function (Bannar-Martin *et al*., 2018).

The third key result is that other predictors, such as temperature and biomass, were particularly influential in all our analyses. That is, the grain dependence of the relationship between S and NPP was coupled with a clear increase in the combined effect of annual temperature and precipitation on both S and NPP towards coarse grains, which supports the notion that either temperature-dependent diversification (Rohde, 1992; Allen *et al*., 2002), niche conservatism (Qian & Ricklefs, 2016), or ecological limits (Šímová *et al*., 2011) shape diversity at these spatial grains. The weaker (relatively to temperature) effect of precipitation is expected since we focus on forests, which only grow above certain precipitation thresholds (Whittaker, 1975). The clear importance of temperature, biomass, longitude, and other predictors such as forest age, temperature seasonality, or species pool (Figs. 4 & 6) highlights that even when the NPP→S relationship holds across grains, other drivers are considerably more important in predicting both (e.g., Ratcliffe *et al*., 2017). Hence, integrating the environmental context surrounding ecological communities into modeling diversity-productivity relationships is a necessary step towards making robust predictions of either biodiversity or ecosystem functioning at any spatial grain.

Our results reveal that mechanisms associated with one direction of diversity-productivity relationships may provide insight to observed patterns of either direction, despite being initially formulated at a different spatial grain. For example, the strong effect of the East-West spatial coordinate on both S and NPP at the fine spatial grain (Fig. 6) suggests that biogeographical history may play a role in shaping the diversity and ecosystem functioning of plant communities, which was initially tested at larger spatial grains (e.g., Hawkins *et al*., 2011; Conradi *et al*., 2020). Increasingly, macroecological mechanisms such as speciation gradients (Schluter & Pennell, 2017) and water-energy variables are being examined in small-grain experimental grasslands to explore their role in mediating niche-based processes (Zuppinger-Dingley *et al*., 2014) and biodiversity effects (Wagg *et al*., 2017), respectively. Similarly, efforts to upscale biodiversity effects on productivity - developed initially to identify local scale mechanisms (Loreau & Hector, 2001; Turnbull *et al*., 2016) - may identify new mechanisms that underpin spatial variation in ecosystem functioning at large spatial scales (Gonzalez *et al*., 2020). An emerging challenge to these efforts is the creation of data products that capture similar processes across spatial scales and are independent (Supplemental Note 2 and Table S3); many of the variables used in this study share similar data sources (e.g. MODIS and LANDSAT sensors), but are ultimately derived from different types of intermediate products. Rather than uniquely focusing on the direction and strength of S-NPP once accounting for other factors, our results show that mechanisms associated with S→NPP and NPP→S likely underpin the context dependency of diversity-productivity relationships across spatial grains (Table 1). These recent developments in BEF research and macroecology suggest that conceptual integration between these two disciplines is just beginning (Craven *et al*., 2019), yet further efforts to bridge disciplinary gaps are essential to deepen current understanding of mechanisms that underpin the shifts in diversity-productivity relationships across spatial scales.

To conclude, we show that the relationship between diversity and productivity strengthens toward coarse grains. This result is in line with expectations from both BEF theory, and some (but not all) expectations from macroecological studies on NPP→S, and highlights the potential of demographic stochasticity and sampling effects to distort or mask diversity-productivity relationships at fine grains. Moreover, we find similar support for both directions of diversity-productivity relationships across spatial grains, revealing that biodiversity and productivity can be both cause and effect. Future research on this relationship needs to move from fine-grain experiments and observational studies to coarse grains in order to fully understand and predict the impacts of anthropogenic biodiversity change on ecosystem function.

## Acknowledgments

All authors recognize support from the German Centre for Integrative Biodiversity Research (iDiv) Halle-Jena-Leipzig (DFG-FZT 118). MvdS is supported by the Rubicon research programme with project number 019.171LW.023, which is financed by the Netherlands Organisation for Scientific Research (NWO). CM acknowledges funding from the Volkswagen Foundation through a Freigeist Fellowship. We thank John Kartesz and Misako Nishino for generously providing access to BONAP data. We thank David Currie and Antonin Machac for initial discussions and Christian Wirth, Katie Barry, Nico Eisenhauer, Stan Harpole, Miguel Mahecha, and the CAFE discussion group for their suggestions to improve analyses.

## Biosketch

The authors are a group of (mostly) early career researchers united by their interest in ecological synthesis, in areas ranging from macroecology to experimental ecology, ecological theory, and ecological modeling.

