## Supplementary Information for "A cross-scale assessment of productivity-diversity relationships"

This file includes:

Supplementary note 1 &2
Figures S1 to S12
Tables S1 to S3

**Supplementary Note 1: Classification of species as forest or non-forest species**

We excluded species that are likely non-forest species based on species occurrences within forested pixels recorded in FIA plots or as occurrence records in GBIF. To this end, we first measured the precision of the coordinates of all GBIF and FIA occurrences. We then extracted raster values from both our forest biomass layer and the forest extent layer (see 'Methods' in main text). We then assigned different levels of certainty to whether the indicated coordinates are located in forest:

- 'high-precision, high-forest': coordinates with ≥ 0.01° precision (ca. 1 km at the equator), and either some forest biomass (>0) and/or forest cover >50% in the pixel.
- 'low-precision, high-forest': coordinates with = 0.1° precision (ca. 11 km at the equator), and either some forest biomass (>0) and/or indicated forest cover >50% in the pixel.
- 'high-precision, low-forest': coordinates with ≥ 0.01° precision, and either some forest biomass (>0) and/or forest cover >20% but <50% in the pixel.
- 'low-precision, low-forest': coordinates with = 0.1° precision, and either some forest biomass (>0) and/or forest cover >20% but <50% in the pixel.

Coordinates falling in pixels for which neither any forest biomass nor forest cover >20% was indicated were classified as 'non-forest'. For each species, we counted the number of total occurrences and the numbers of occurrences falling in each level of certainty. We calculated a weighted sum of forest occurrences for each species, weighting 'high-precision, high-forest' occurrences by 100%, 'low-precision, high-forest' occurrences by 50%, 'high-precision, low-forest' occurrences by 20%, and 'low-precision, low-forest' occurrences 5%. We defined species as likely to occur in forests if they had either ≥ 5 weighted forest occurrences or ≥ 1/5 of all their weighted occurrences in forests.

**Supplementary Note 2: Potential circularity in environmental predictors**

Many of the environmental predictors that we use were generated using complex algorithms that may potentially rely on similar types of primary data or geographic layers. Examples of these common data sources are interpolated climatic layers, layers derived from MODIS and LANDSAT sensors, or the use of digital elevation models (DEMs) for interpolation purposes (Table S3). Thus, environmental variables derived from similar data are expected to be correlated. This can lead to circularity when their relationship is assessed in statistical analyses, since the same or similar data have been used to create both the response and the predictor. Here we briefly describe how this may apply to our analyses, and why we still consider our results valid.

**Algorithms and ingredients differ.** Table S3 shows that although many variables ultimately share similar data sources, e.g. the same remote sensor or weather stations, they are actually based on different types of intermediate products, and these are always combined with a unique combination of other data. For example, the MODIS-derived NPP is based on the MOD15 product (fraction of photosynthetically active radiation) and its combination with two local minimum and maximum temperature and vapor pressure deficit (Zhao & Running 2010, and see also ). In contrast, even though the biomass layer (Blackard et al. 2007) also uses MODIS data, it is based on a different set of layers, i.e. percentage tree cover, surface reflectance, and vegetation indices. The climatic data that the biomass layer uses are also different, which in this case are long term averages. This leads to imperfect correlations between these variables even at the coarsest grain (Fig. S7), which still leaves space for their independent effects on richness, or NPP.

**S-NPP relationships.** Another reason why our results are valid is our focus on S-NPP relationships. We estimated S at the fine spatial grain using FIA data (USDA Forest Service, 2017), and estimated S at the intermediate and coarse spatial grains using data from BONAP (Kartesz, 2015). In both cases, the data used for estimating S are completely independent of the data used for estimating NPP.

**Our results hold when using plot-derived NPP.** In Figure S12 we show that the S-NPP relationship remains weak, relative to other predictors, regardless of whether MODIS-derived or plot-derived NPP data are used, where the latter is unaffected by the circularity issue. However, this only applies to the fine spatial grain data.

**Our results reveal weak S-NPP relationships across spatial grains.** This further supports the notion that data circularity does not strongly influence our results, since we expect that it should lead to strong, rather than weak, S-NPP relationships.

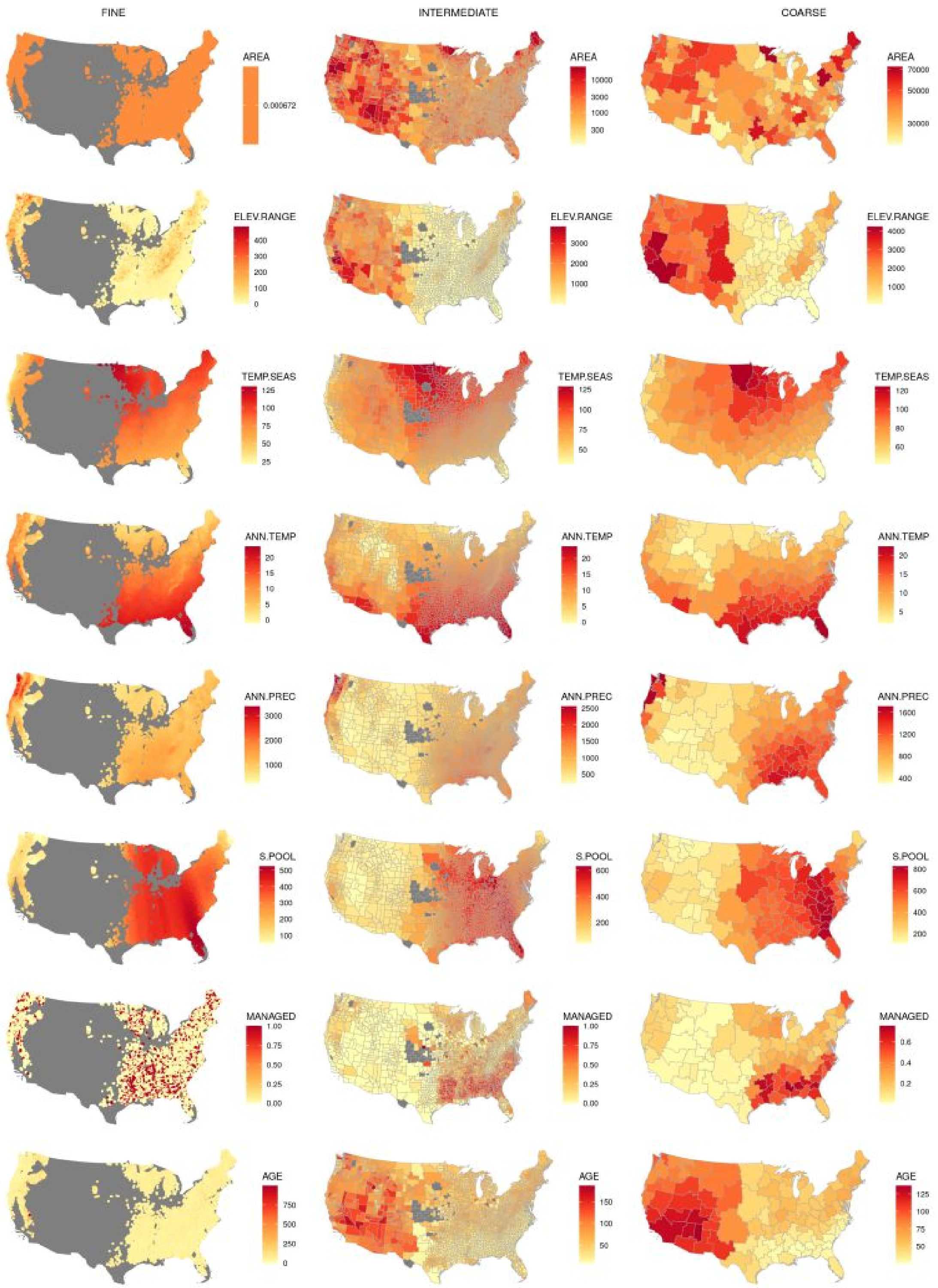


**Fig. S1.** Maps of all of the covariates at FINE (left column), INTERMEDIATE (middle column) and COARSE (right column) spatial grains.

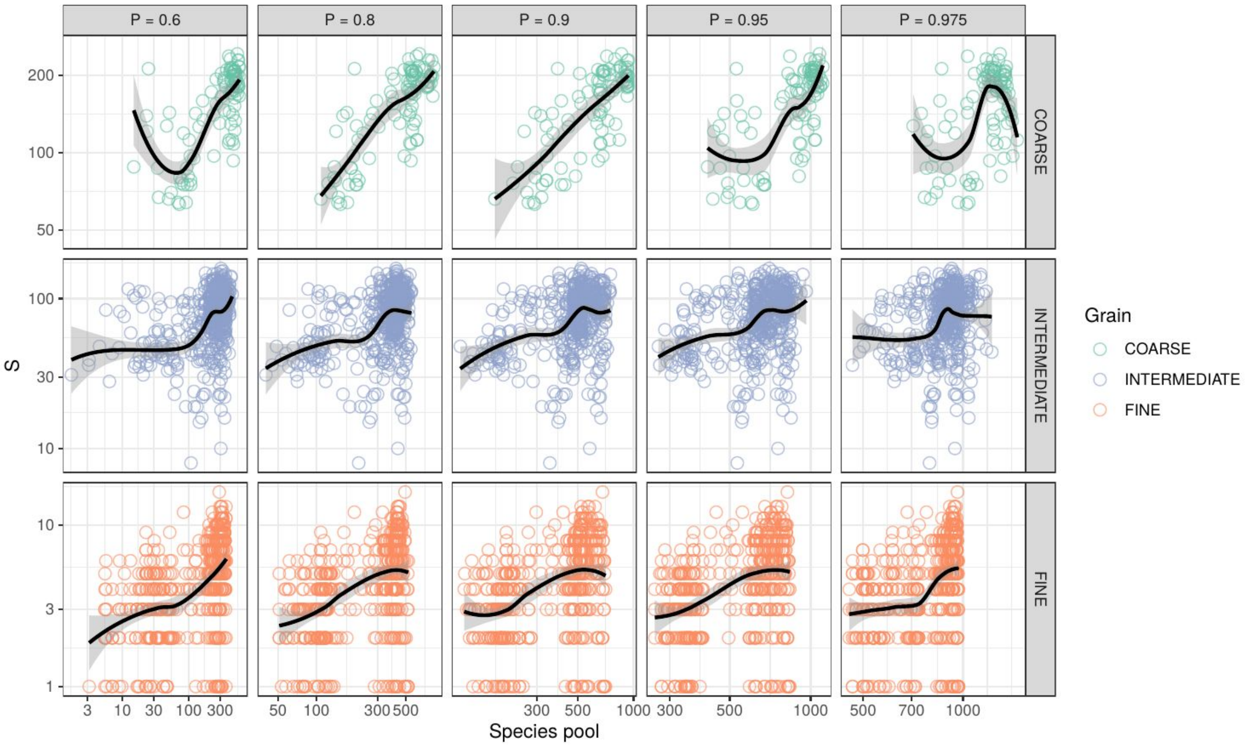


**Fig. S2**. Relationships between species richness (S) and five types of species pools derived using five alternative exponential distance-decay functions, with scaling coefficients *P* that determined the probability of a species occurring in neighboring units would disperse to the focal unit of 0.975, 0.95, 0.90, 0.80, and 0.60 (columns). For the main analysis, we chose *P* = 0.8 as it correlates with S at all three grains. Solid lines are smoothing splines, grey shading indicates standard errors.

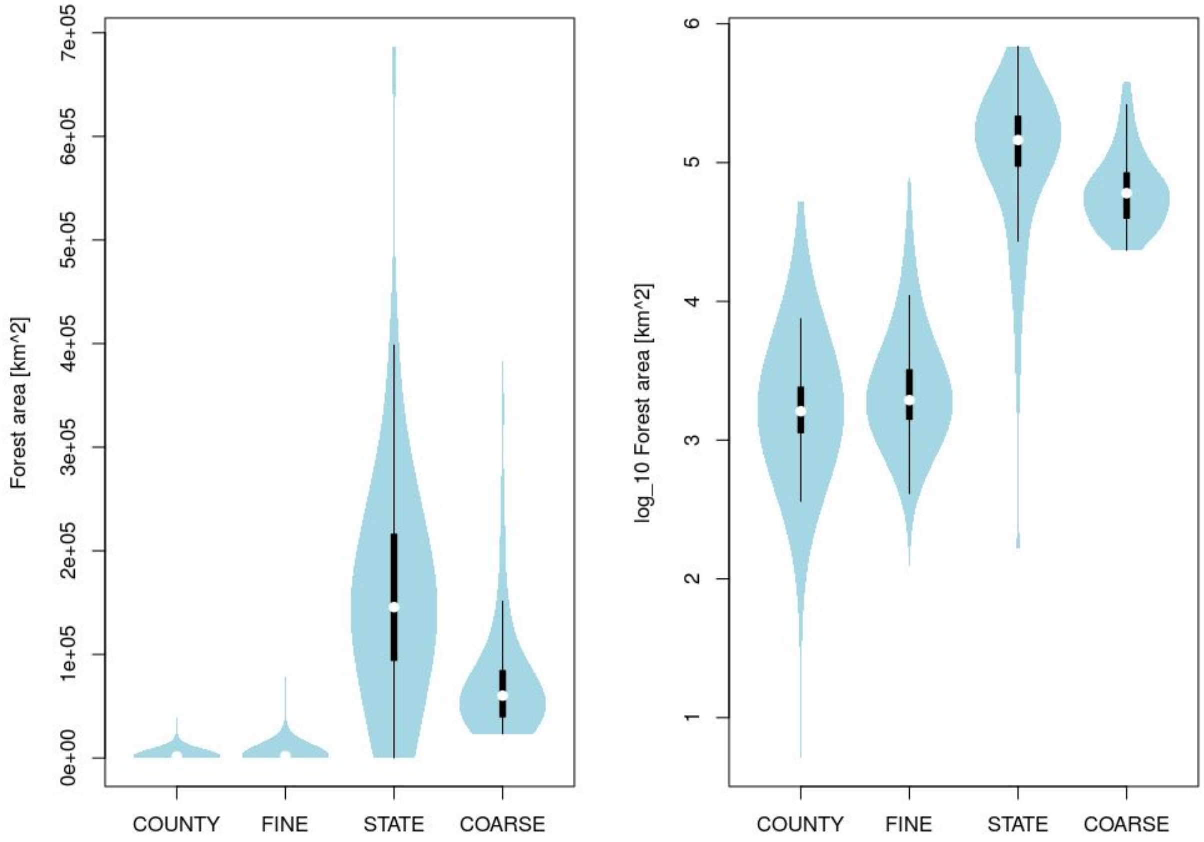


**Fig. S3.** The effect of using the spatial aggregation algorithm (see Methods) on the variation of forest area within the spatial units. COUNTY and STATE are the two original spatial grains of the raw species richness data (BONAP; Kartesz, 2015), which represent US administrative counties and states respectively. The alternative spatial grains that we use in our analysis are FINE and COARSE, which were created by incremental aggregation of the COUNTY data, while minimizing the variation of forest area. Left panel uses linear scale, right panel uses $log_{10}$ scale.

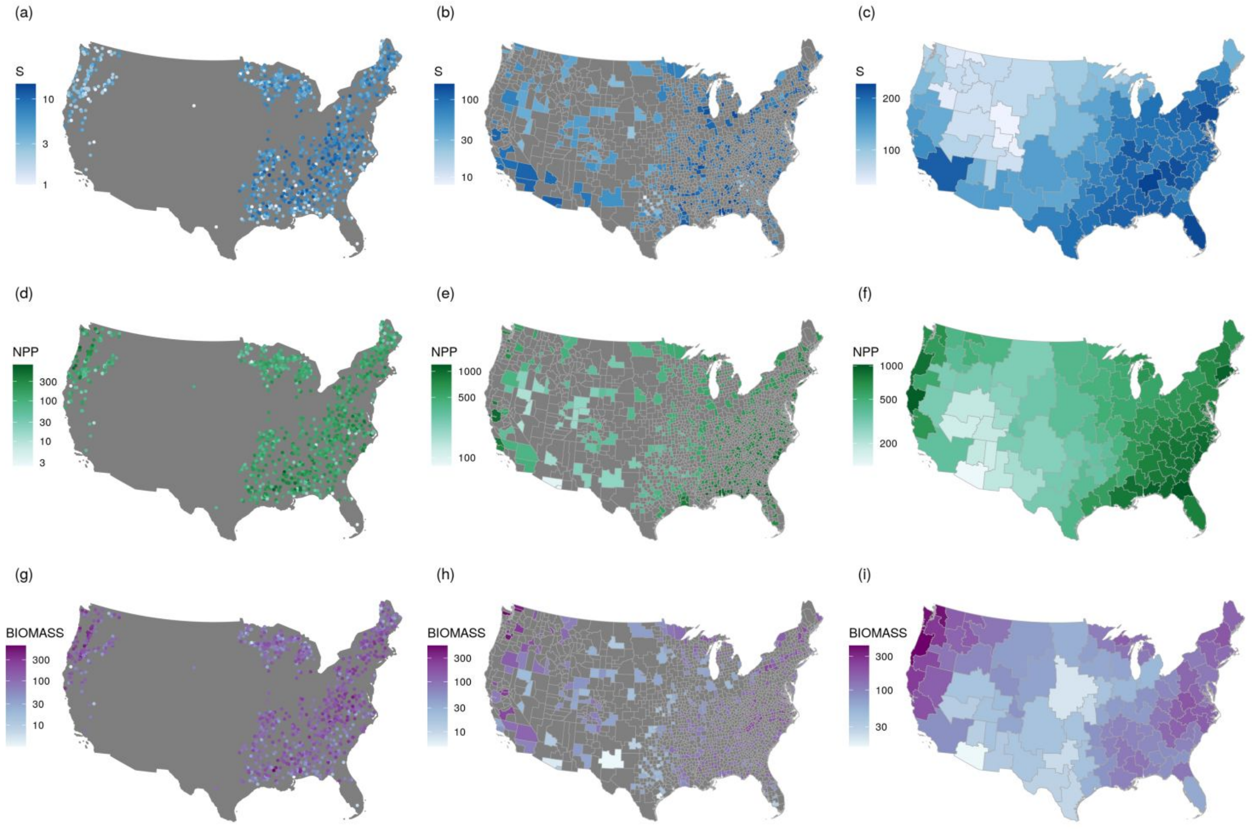


**Fig. S4.** Maps of the *subsampled* focal variables: S, NPP and BIOMASS at three spatial grains. Maps at the FINE (a,d, g) and INTERMEDIATE (b, e, h) spatial grains show stratified random samples of 1,000 and 500 locations respectively. The values in all plots use $\log_{10}$ scale.

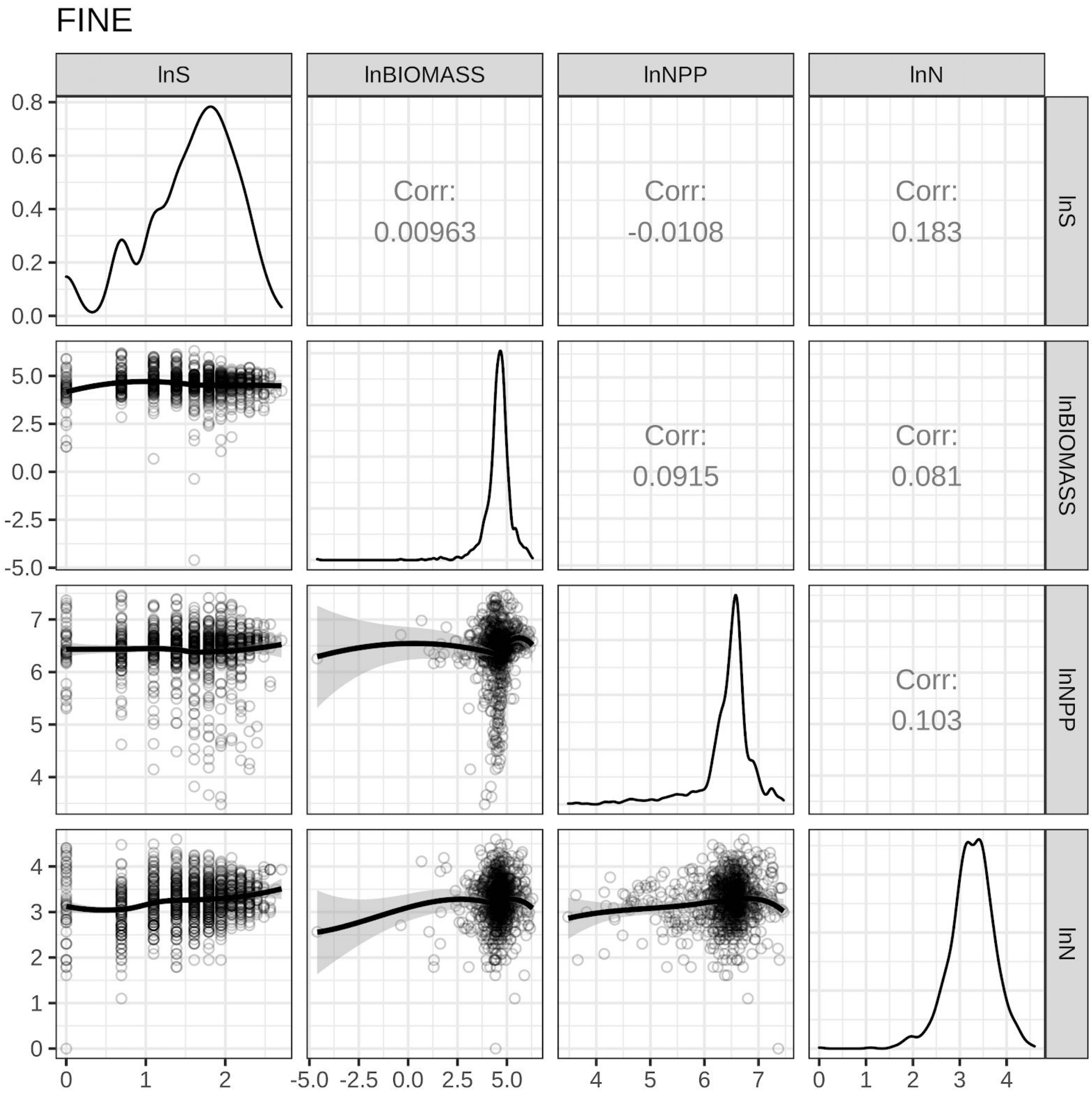


**Fig. S5.** Pairwise relationships between the MODIS-derived NPP, S and biomass data at the fine spatial grain (N = 1,000). All variables were natural-log transformed.

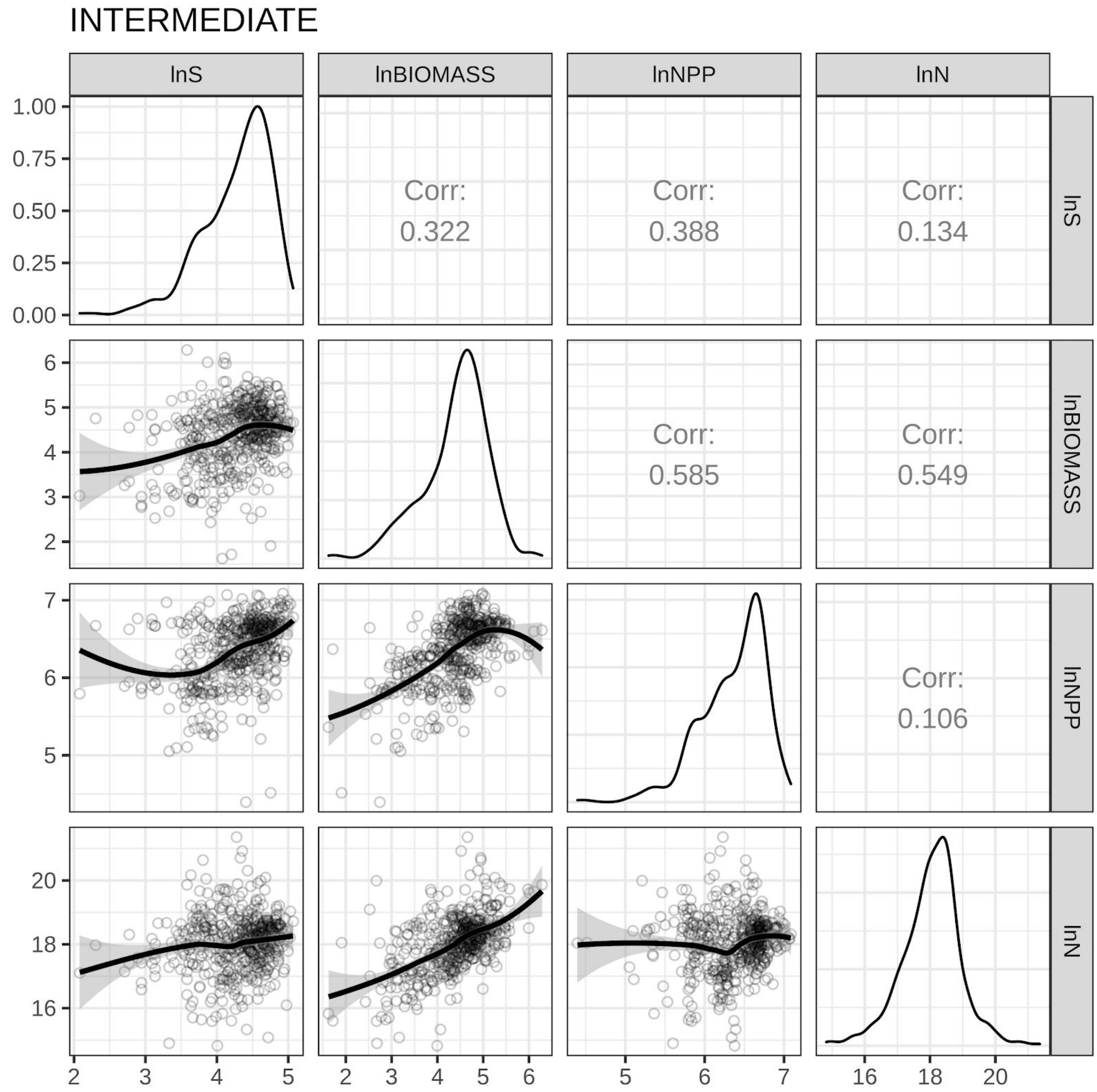


**Fig. S6.** Pairwise relationships between the MODIS-derived NPP, S and biomass data at the intermediate spatial grain (N = 500). All variables were natural-log transformed.

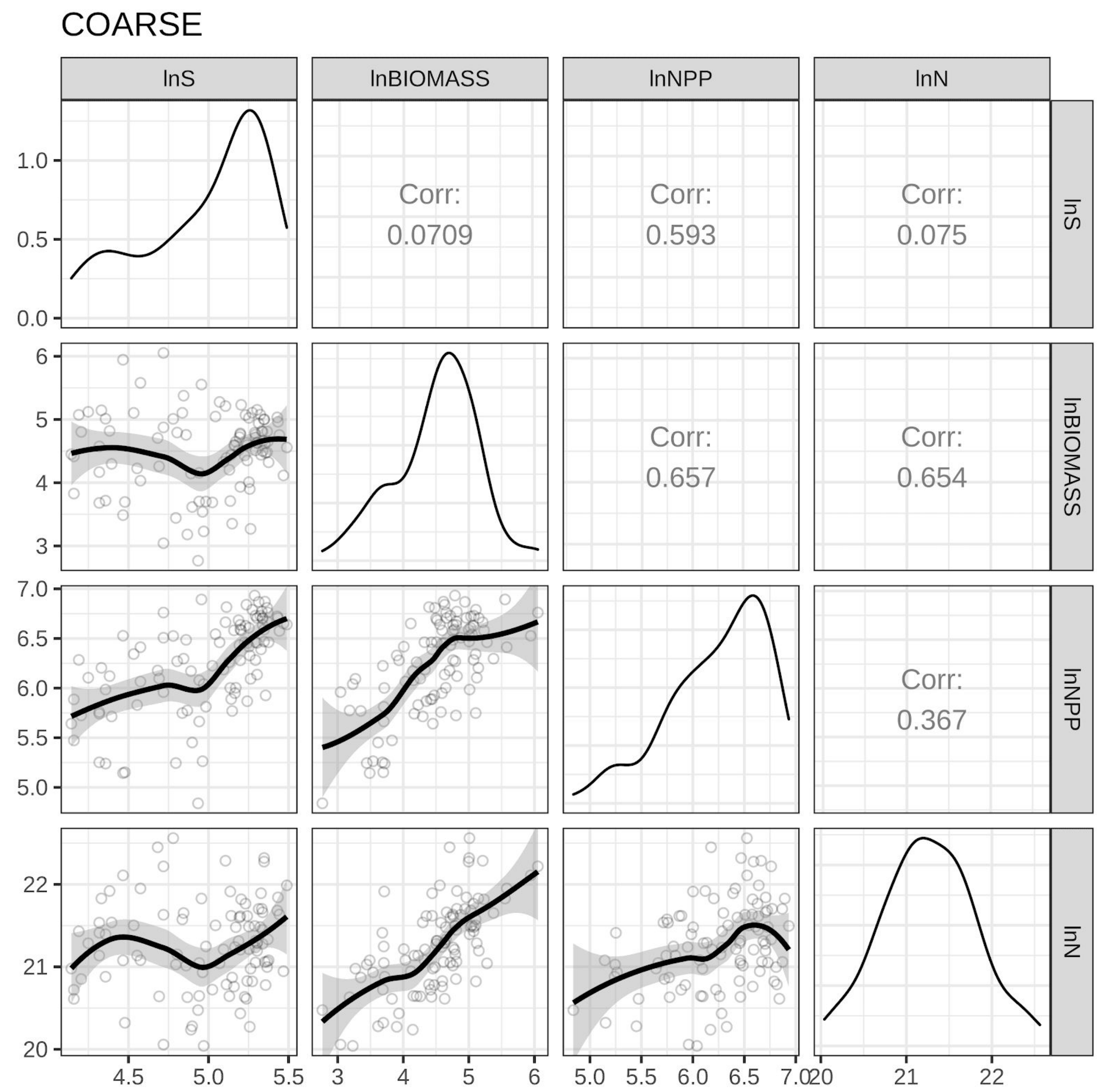


**Fig. S7.** Pairwise relationships between the MODIS-derived NPP, S and biomass data at the intermediate spatial grain (N = 98). All variables were natural-log transformed.

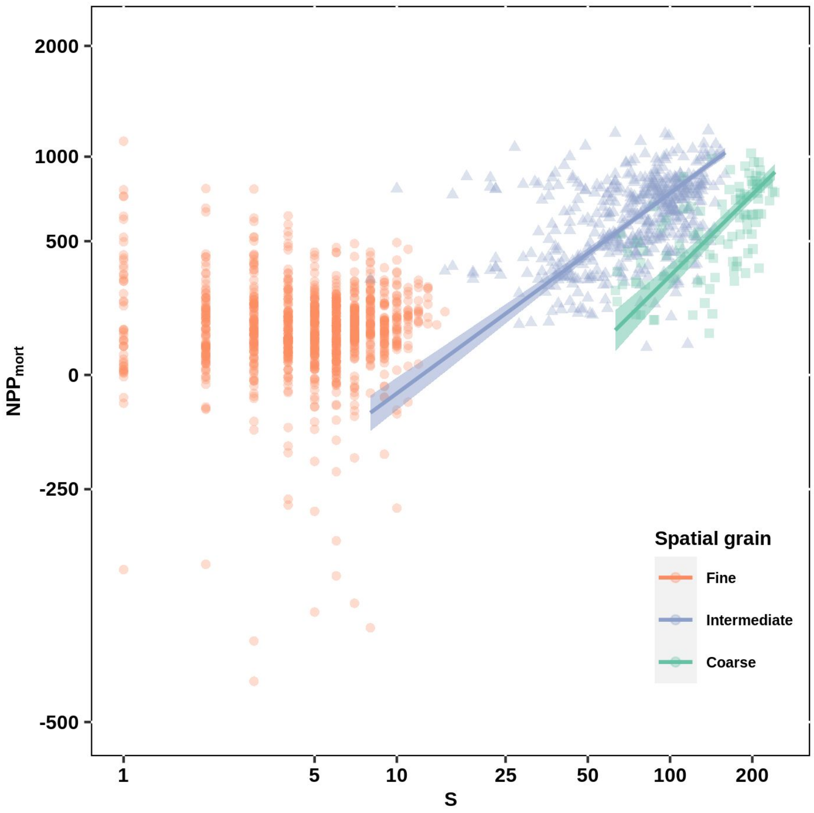


**Fig. S8.** Bivariate relationships between observed species richness (S) and productivity ($NPP_{mort}$) of forests at three spatial grains across the contiguous USA. NPP is estimated as the sum of aboveground C growth of living trees, ingrowth by recruitment, and loss from tree mortality. NPP is MODIS-derived at the intermediate and coarse spatial grains. Solid lines are standardised major-axis regressions fitted at each spatial grain and shaded areas are 95% confidence intervals; only regressions with statistically significant slopes ($P<0.05$) were visualised. Note that axes are on the natural log scale. Analyses were performed using stratified random samples of 1000, 500 and 98 spatial units at the fine, intermediate and coarse spatial grains, respectively.

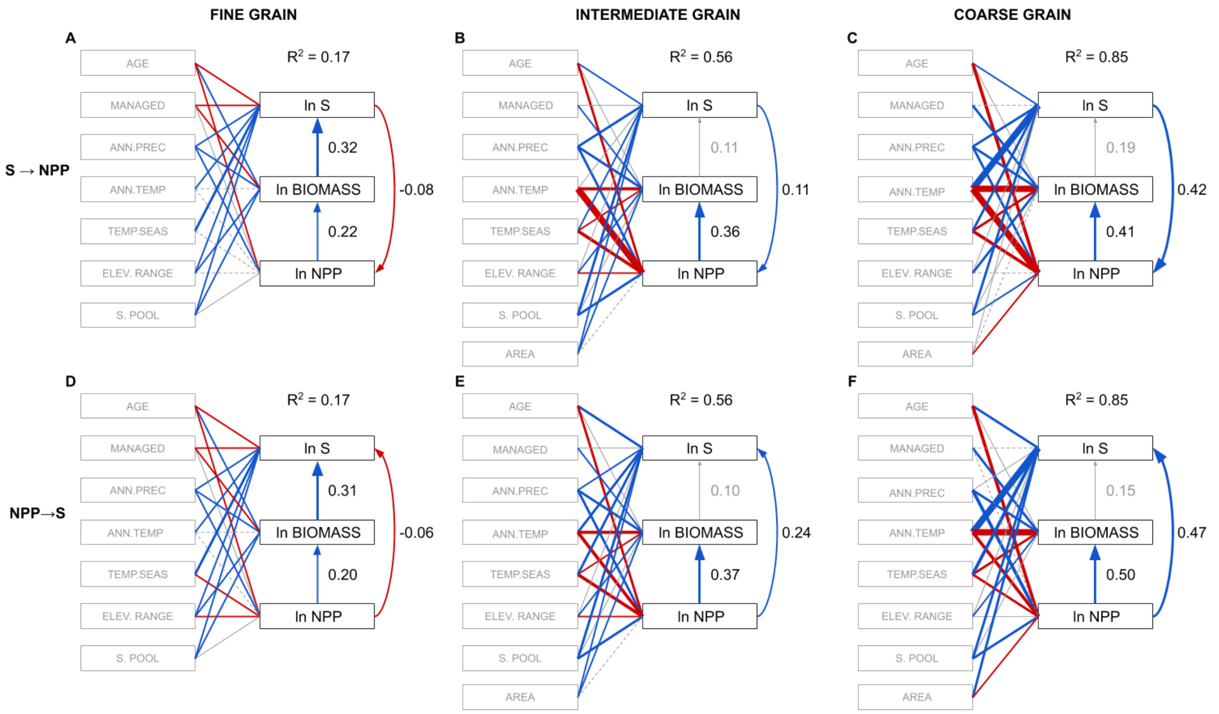


**Fig. S9.** Structural equation models (SEM) testing the influence of diversity (S) on productivity (NPP)('S → NPP'; A, B, C) and that of NPP on S ('NPP → S'; D, E, F), once controlling for environmental variables (e.g., mean annual precipitation, mean annual temperature, temperature seasonality, and elevation range), size of the species pool, forest age, and management, in forests across the contiguous USA at three spatial grains. All models fit the data well at all spatial grains (P-value of the Chi-square test > 0.1; Table S2). Boxes represent measured variables and arrows represent relationships among variables. Solid blue and red arrows represent significant (P< 0.05) positive and negative standardized path coefficients, respectively, and their width is scaled by the corresponding standardized path coefficient. Solid and dashed gray arrows represent non-significant (P>0.05) positive and negative standardized path coefficients, respectively. $R^2$ is the average of $R^2$ values for S, BIOMASS, and NPP. NPP is estimated using plot-level data (NPP.mort) at the fine spatial grain and using MODIS-derived data at intermediate and coarse spatial grains. AGE is forest age, MANAGED is forest management, ANN.PREC is mean annual precipitation, ANN.TEMP is mean annual temperature, TEMP.SEAS is temperature seasonality, ELEV.RANGE is elevation range, S.POOL is the regional species pool, and AREA is area. S, BIOMASS, NPP (MODIS-derived), and AREA were natural log transformed prior to analysis.

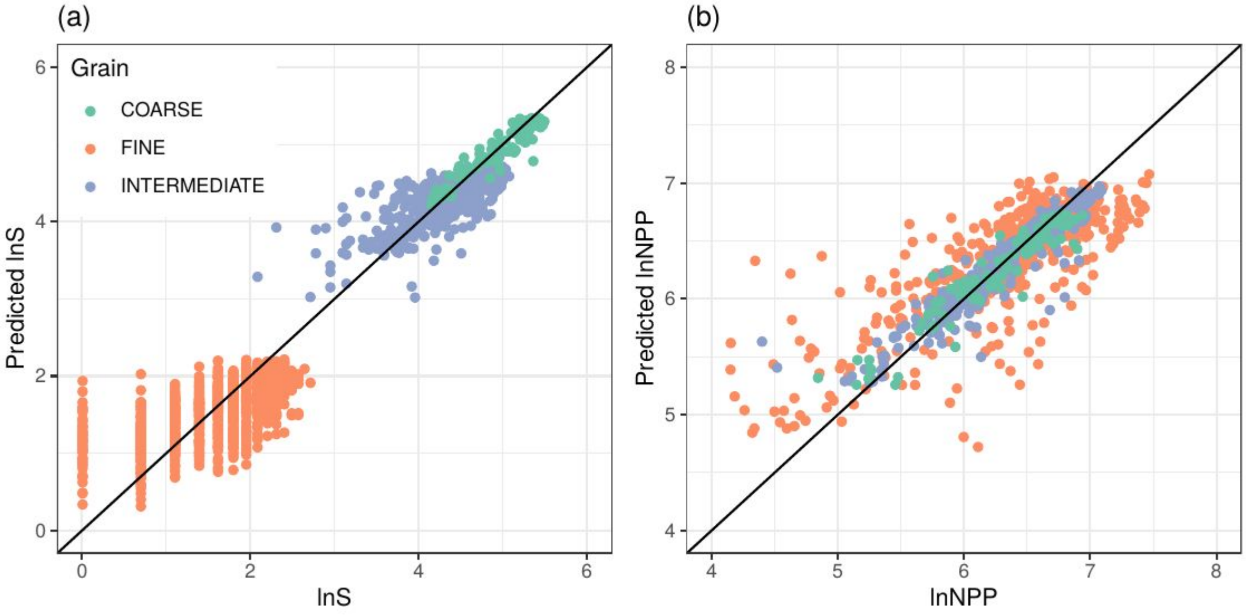


**Fig. S10.** Observed versus predicted values from the random forest models at three spatial grains. NPP is MODIS-derived at all grains.

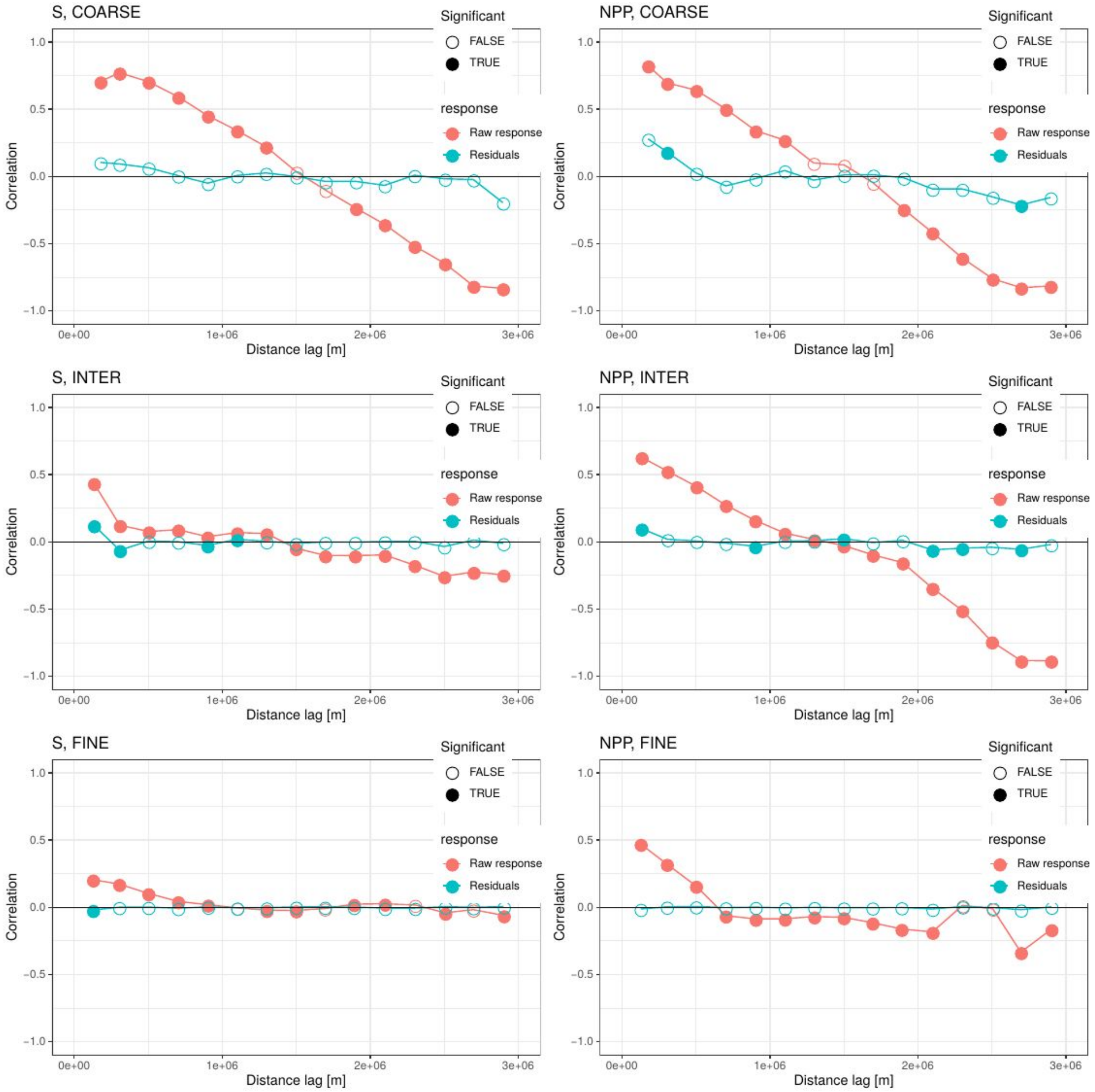


**Fig. S11.** Spatial correlograms of species richness (S, left column) and MODIS-derived NPP (right column), both in their raw form (red) and as residuals from the random forest models (blue), assessed at three spatial grains (rows). Although raw S and NPP are significantly correlated at all distance lags up to 1,000 km (1e+06 m), spatial correlation in the model residuals is negligible. This is achieved by using the X and Y spatial coordinates in the random forest models.

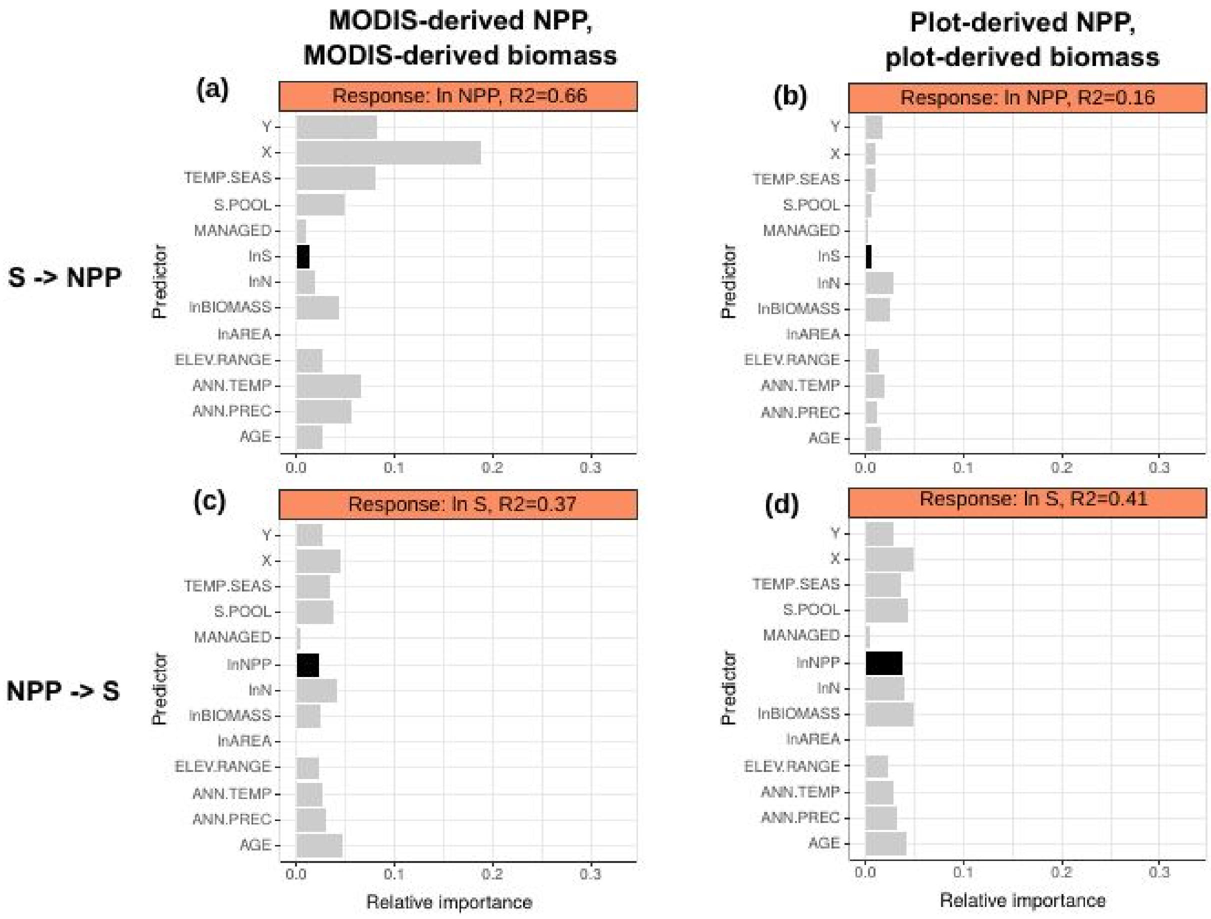


**Fig. S12.** Comparison of random forest models that use different measures of NPP and biomass at the FINE spatial grain. For (a) and (c), NPP is estimated using MODIS, while for for (b) and (d) it is directly estimated using plot data and as the sum of aboveground C growth of living trees, ingrowth by recruitment, and loss from tree mortality.

**Table S1.** Overview of all variables used in the SEM and RFs.

| Variable | Abbreviation | Units | The grain at which data are used | Source, reference | Layer resolution, if applicable |
|---|---|---|---|---|---|
| Species richness | S | # of species | fine | FIA plots (USDA Forest Service, 2017) | |
| | | | intermediate, coarse | BONAP (Kartesz, 2015) | |
| Field net primary productivity, with mortality | $NPP_{mort}$ | $gC/m^2/year$ | fine | FIA plots (USDA Forest Service, 2017) | |
| MODIS-derived net primary productivity | NPP | $gC/m^2/year$ | all grains | MODIS GPP/NPP MOD17 project (Zhao *et al.* 2005, Zhao & Running 2010) | 1 $km^2$ |
| Biomass | BIOMASS | Mg/ha | fine | FIA plots (USDA Forest Service, 2017) | |
| | | Mg/ha | fine, intermediate, coarse | Aboveground live forest biomass map (Blackard *et al.* 2008) | 1 $km^2$ |
| Number of trees | N | # of trees | fine | FIA plots (USDA Forest Service, 2017) | |
| | | # of trees | intermediate, coarse | Global tree density map (Crowther et al. (2015) | 1 $km^2$ |
| Forest age | AGE | years | fine | FIA plots (USDA Forest Service, 2017) | |
| | | years | intermediate, coarse | NACP (Pan et al. 2012) | 1 $km^2$ |
| Management regime | MGMT | managed or not | fine | FIA plots (USDA Forest Service, 2017) | |
| | | proportion of managed FIA plots | intermediate, coarse | FIA plots (USDA Forest Service, 2017) | |
| Mean annual precipitation | MAP | average mm/y | all grains | WorldClim 1.4 (Hijmans et al. 2005) | 1 $km^2$ |
| Mean annual temperature | MAT | average ℃/y | all grains | WorldClim 1.4 (Hijmans et al. 2005) | 1 $km^2$ |
| Temperature seasonality | TEMP.SEAS | standard deviation of monthly ℃ * 100 | all grains | WorldClim 1.4 (Hijmans et al. 2005) | 1 $km^2$ |
| Elevation range (altitudinal span) | ELEV.RANGE | m | all grains | SRTM v 2.1 (USGS 2009) | 30 arc-sec |
| Size of species pool | S.POOL | # of species | all grains | FIA, GBIF, BONAP (see Methods) | |

**Table S2.** Model fit of structural equation models (SEM) testing the influence of diversity on productivity and that of NPP on S, once controlling for environmental variables, size of the regional species pool, forest age, and management in forests across the contiguous USA at three spatial grains. Model fit to the data was accepted if the *P* of the $\chi^2$ statistic test was greater than >0.05. Degrees of freedom correspond to that of the $\chi^2$ statistic test for each model and spatial grain. Please note that S → NPPmort and NPPmort → S were only fit at the fine spatial grain, as MODIS-derived NPP was used at the intermediate and spatial grains.

| **Model** | **Spatial grain** | ***Degrees of freedom*** | $\chi 2$ | ***P*** |
|---|---|---|---|---|
| S → NPP | fine | 1 | 0.39 | 0.53 |
| S → NPP | intermediate | 1 | 0.13 | 0.72 |
| S → NPP | coarse | 1 | 0.04 | 0.84 |
| NPP → S | fine | 1 | 1.19 | 0.28 |
| NPP → S | intermediate | 1 | 0.05 | 0.82 |
| NPP → S | coarse | 1 | 0.75 | 0.39 |
| S → NPPmort | fine | 1 | 0.07 | 0.80 |
| NPPmort → S | fine | 1 | 0.04 | 0.84 |

**Table S3.** Variables that have been derived from similar data sources.

| | **MODIS-derived NPP** | **Biomass** | **N** | **Forest Age** | **WorldClim** |
|---|---|---|---|---|---|
| **Reference** | Zhao et al. (2005), Zhao & Running (2010) | Blackard et al. (2007) | Crowther et al. (2015) | Pan et al. (2012) | Hijmans et al. (2005) |
| **Validation** | EMDI data of local NPP measures | Biomass from FIA dataset | A collection of forestry surveys across the world | Forest age from FIA dataset | A collection of datasets across the world |
| **Interpolated climatic data from weather stations** | GMAO/NASA interpolated data of min and max temperature and vapor pressure deficit | PRISM data for temperature and precipitation | yes (details not specified) | no | A collection of datasets across the world |
| **MODIS** | MOD15 (fraction of photosynth active radiation, FPAR) | MOD44 (percent tree cover), MOD09 (surface reflectance), MOD13 (veg. index composites) | Enhanced vegetation index (EVI, details not specified, likely MOD13) | no | no |
| **LANDSAT** | no | land cover classes | no | LEDAPS fractions of disturbed areas | no |
| **Digital elevation model (DEM)** | May appear in interpolation of the climatic data | yes | yes | no | Appears in interpolation of the climatic data |
